# Loss of m^1^acp^3^Ψ ribosomal RNA modification is a major feature of cancer

**DOI:** 10.1101/840132

**Authors:** Artem Babaian, Katharina Rothe, Dylan Girodat, Igor Minia, Sara Djondovic, Miha Milek, Sandra E. Spencer Miko, Hans-Joachim Wieden, Markus Landthaler, Gregg Morin, Dixie L. Mager

## Abstract

The ribosome is an RNA-protein complex essential for translation in all domains of life. The structural and catalytic core of the ribosome is its ribosomal RNA (rRNA). While mutations in ribosomal protein (RP) genes are known drivers of oncogenesis, oncogenic rRNA variants have remained elusive. We discovered a cancer-specific single nucleotide variation in 18S rRNA at nucleotide 1248.U in up to 45.9% of colorectal carcinoma (CRC) patients and present across >22 cancer types. This is the site of a unique hyper-modified base, 1-methyl-3-α-amino-α-carboxyl-propyl pseudouridine (m^1^acp^3^Ψ), a >1 billion years conserved RNA modification at the ribosome’s peptidyl decoding-site. A sub-set of CRC tumors we term ‘hypo-m^1^acp^3^Ψ’, show sub-stoichiometric m^1^acp^3^Ψ-modification unlike normal control tissues. A m^1^acp^3^Ψ knockout model and hypo-m^1^acp^3^Ψ patient tumors share a translational signature, characterized by highly abundant ribosomal proteins. Thus, m^1^acp^3^Ψ-deficient rRNA forms an uncharacterized class of ‘onco-ribosome’ which may serve as a chemotherapeutic target for treating cancer patients.

## Introduction

The ribosome is a massive ribonucleoprotein particle (RNP) responsible for the transformation of genetic information encoded as nucleic acids into functional proteins encoded as amino acids. Unlike most RNPs, it is ribosomal RNA (rRNA), and not ribosomal proteins (RPs) that form the most ancient and catalytic core of the complex (Cech, 2000). rRNA is further functionalized by a constellation of at least 14 distinct chemical modifications across 200+ sites, (Taoka et al., 2016) clustering around active sites of the ribosome (Sloan et al., 2017), yet the function of many rRNA modifications remains unclear.

The human ribosome contains >80 RPs and four rRNAs, totaling ∼80% of cellular RNA. During the initial human genome sequencing project, ribosomal DNA (rDNA) loci were systematically excluded from the reference genome (Lander et al., 2001); given that a reference sequence of the rRNA gene, *RNA45S*, was available and the 80-800 rDNA copies were believed to be homogeneous (Elder and Turner, 1995). Although there was early evidence for rDNA polymorphism in humans (Kuo et al., 1996; Leffers and Andersen, 1993). Thus, as technology and sequencing consortium projects revolutionized genomics and transcriptomics, our understanding of rDNA variation has lagged. rDNA sequence variation at the intra- and inter-individual level has been documented in multiple species including humans (Bik et al., 2013; Rabanal et al., 2017; Babaian, 2017; Kim et al., 2018; Parks et al., 2018), but the functional implications of rDNA variation remain elusive. Mutation of RP genes and ribosome biogenesis factors can cause a class of diseases termed ribosomopathies, including Diamond Blackfan anemia (DBA) (Nakhoul et al., 2014), and some cancers (Gourdazi and Lindstrom). It has been hypothesized that cancer cells contain a functionally specialized class of ribosomes to facilitate rapid protein synthesis, termed the “onco-ribosomes” (Shi and Barna, 2015; Dinman, 2016). Cancer genomics has supported this notion with the identification of several oncogenic driver mutations in RP genes (Gourdazi and Lindstrom, 2016), the best characterized of which are RPL10 (uL16) p.R98S in T-cell acute lymphoblastic leukemia (Girardi et al., 2018; Kampen et al., 2019) and RPS15 (uS19) C’ terminal mutations in chronic lymphocytic leukemia (Bretones et al., 2018). In addition, germline mutations such as in DBA patients and in *RPS20* can cause heredity cancers including colorectal carcinoma (CRC) (Vlachos et al., 2012; Nieminen et al., 2014).

As RP mutations have been implicated in tumorigenesis, we hypothesized that rRNA variation or mutation is a cancer driver. To map functional rRNA sequence variation, we considered tumorigenesis as a natural experiment in which polymorphic and mutant rRNA alleles undergo selective evolutionary change in frequency within each patient. We discovered a surprising 18S rRNA single nucleotide alteration at the decoding core of the ribosomal peptidyl (P)-site, affecting up to 45.9% of CRC patients, making this the most frequent ribosomal variant associated with cancer to date and potentially revolutionizing future chemotherapeutic strategies against this disease.

## Results & Discussion

### An unexpected rRNA variant in cancer: sub-stoichiometric modification of 18S.1248.m^1^acp^3^Ψ

In an initial screen for cancer-driver rRNA variants, we aligned RNA-seq reads from 66 colorectal carcinoma (CRC) tumors and patient-matched adjacent normal tissue to a single-copy reference rDNA. To test for allelic selection inconsistent with neutral drift, the patient-matched difference in expressed variant allele frequency (VAF) was measured for deviation from zero for each position of 18S and 28S (Fig. 1a). Non-reference reads at a single nucleotide deviated from neutrality, 18S:r.1248.U (p_adj_ = 3.81e-8). In this cohort, the 18S.1248.U reference allele is recurrently selected over non-U or 18S.1248V alleles in a striking 44.9% of CRC patients (Fig. 1a); in comparison, oncogenic *KRAS*^G12^ codon mutation occurs in only 36% of CRC patients (Tan and Du, 2012).

**Figure 1:**
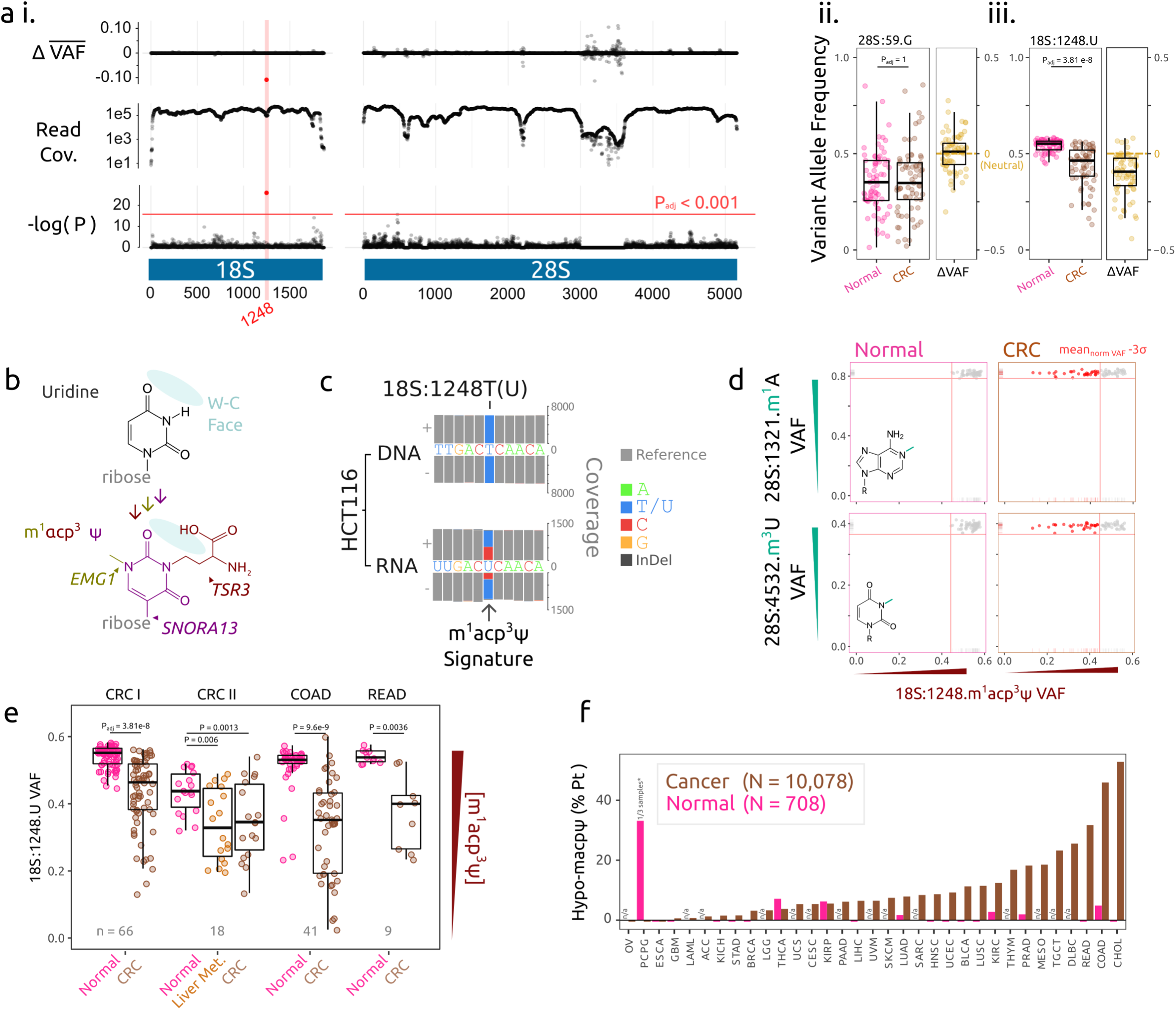
The hypo-m^1^acp^3^Ψ phenotype in cancer. **a i**, Screen for change in the average variant allele frequency (VAF) across *18S* and *28S* ribosomal RNA (rRNA) in colorectal cancer (CRC) RNA-seq compared to patient-matched normal epithelium controls (n = 69). Read coverage and quality drops at extreme GC-content (>90%) regions of 28S, these low-coverage regions were excluded from further analysis. **ii**, The common human rRNA polymorphism 28S:r.59G>A ranges from 0.05–0.93 DNA allele frequency (Babaian, 2017) and was expressed comparably in the normal epithelium between variant allele frequencies (VAF) of 0.01-0.86. Neither allele is directionally selected for during cancer evolution, consistent with neutral drift (p_adj_ = 1, t = -0.44). **iii**, 18S:r.1248.U is significantly enriched (p_adj_ = 3.81e-8, t = 8.33) for the reference U allele. **b**, The 18S:r.1248.U base normally undergoes enzymatic hyper-modification to 1-methyl-3-α-amino-α-carboxyl-propyl pseudouridine (m^1^acp^3^Ψ) in three steps; *SNORA13* guided pseudouridylation; EMG1 N1-methylation; and finally 3-amino-carboxyl-propylation by TSR3, which are not down-regulated in hypo-m^1^acp^3^Ψ tumors (Fig. S1d). **c**, Perturbation of the Watson-Crick face of the modified base results in a distinct nucleotide misincorporation signature by reverse transcriptase in first strand cDNA synthesis, which is read out on both the first (+) and second read-pair (-) of sequencing of matched HCT116 whole genome DNA- and RNA-seq (Cancer Cell Line Encyclopedia, (Ghandi et al., 2019)). **d**, Patient 18S:r.1248.m^1^acp^3^Ψ hypo-modification is defined as a decrease in VAF by three standard deviations (3σ) of the matched-normal samples. Hypo-m^1^acp^3^Ψ is not correlated with the loss of other rRNA modifications detectable by RNA-seq. **e**, The hypo-m^1^acp^3^Ψ phenotype is replicated in three additional, independent cohorts of CRC with patient-matched adjacent normal controls, including two cohorts from The Cancer Genome Atlas (TCGA), colorectal adenocarcinoma (COAD) and rectal adenocarcinoma (READ). **f**, Hypo-modification of 18S:r.1248.m^1^acp^3^Ψ is prevalent but not ubiquitous across the TCGA cancer cohorts (n = 10,078 patients) and largely absent from patient-matched normal controls (n = 708) (see: Fig. S1).

Surprisingly, at the <u>DNA</u> level, the respective nucleotide *RNA45S*:4908.T is invariable in humans and CRC (Babaian, 2017 Fig. S1a,b), and in the mature rRNA this uridine undergoes hyper-modification to 1-methyl-3-α-amino-α-carboxyl-propyl pseudouridine (m^1^acp^3^Ψ) (Fig. 1b) (Taoka et al., 2016). The m^1^acp^3^Ψ modification perturbs standard Watson-Crick base-pairing during cDNA synthesis by reverse transcriptase (RT) (Helm and Motorin, 2017), resulting in base misincorporation and enzyme stalling, which is read-out as a consistent ‘modification signature’ in RNA-seq (Fig. 1c, reviewed in (Helm and Motorin, 2017)). Restated, non-reference or variant reads at 18S.1248.U correspond to m^1^acp^3^Ψ modification and not a genetic variation. Thus, the increase in reference U sequence suggests that the m^1^acp^3^Ψ modification is incomplete or sub-stoichiometric in CRC tumors, which we term the ‘hypo-m^1^acp^3^Ψ phenotype’. The 28S.1321.m^1^A and 28S.4532.m^3^U rRNA modifications also cause a ‘modification signature’ in RNA-seq (Helm and Motorin, 2017). These modifications do not decrease in CRC tumors or matched normal controls, excluding a non-specific rRNA modification effect (Fig. 1d).

The hypo-m^1^acp^3^Ψ phenotype is reproducible at comparable frequency (27.8-45.9%) in three additional independent patient-matched CRC cohorts (Fig. 1e,f). Analysis of 10,036 cancer patients and 712 normal controls across an additional 31 cancer-patient cohorts reveals that hypo-m^1^acp^3^Ψ occurs at a significant frequency across a diverse set of cancers but is not pan-cancerous (global recurrence: 9.6% range: 0-52.8%) (Fig. 1f, S1).

To validate these findings, we designed a simple and rapid <u>a</u>minocarboxyl propyl reverse transcription (aRT)-PCR assay for measuring 18S.1248.m^1^acp^3^Ψ modification. The aRT-PCR is reproducible and quantitative (Fig. S2). A critical technical limitation which also applies to RNA-seq is that different RT enzymes or RT reaction chemistries, differ the base misincorporation rates at 18S.1248.m^1^acp^3^Ψ (Fig. S2d). Thus, cross-cohort or cross-experimental comparisons should be made cautiously (Fig. S2b-d). Indeed, batch-effects on VAF are seen within TCGA cohorts, yet hypo-m^1^acp^3^Ψ replicates across batches further supporting that hypo-m^1^acp^3^Ψ is occurring in tumors specifically (Fig. S1b).

We validated that the hypo-m^1^acp^3^Ψ phenomenon also occurs in CRC cell lines assayed as a single batch and confirmed the results are not a sequencing artifact (Fig. S2e). To test if m^1^acp^3^Ψ-deficient rRNA incorporate into mature ribosomes, we isolated monosomes and polysomes and detected low m^1^acp^3^Ψ modification levels in mono- and di-somes (Fig. S2f).

As the molecular, biological, and medical significance of 18S.1248.m^1^acp^3^Ψ unfolds, it is obvious that genotyping technologies (such as sequencing or our aRT-PCR assay for m^1^acp^3^Ψ) and previous m^1^acp^3^Ψ assays such as primer extension can be adapted as affordable and rapid diagnostic or prognostic assays.

### 18S:1248.m^1^acp^3^Ψ is an ancient modification at the decoding core of the peptidyl-site

We next investigated the evolutionary and structural characteristics of 18S.1248.m^1^acp^3^Ψ for functional insight. The 18S:1248.U base and m^1^acp^3^Ψ modification are absolutely conserved across *Eukarya* at a residue located in the loop of universal helix 31 (Fig. S3). *TSR3* is the aminocarboxylpropyl transferase which deposits the acp_3_ at 18S.1248.U (Meyer et al., 2016) and it only modifies this single rRNA position, in 100% of mature rRNA molecules in *Eukaryotes* (Taoka et al., 2018; Yang et al., 2016).

Structurally, 18S.1248.m^1^acp^3^Ψ is solvent-exposed at the ribosomal P-site, immediately adjacent to the codon:anti-codon interface (Fig. 2). Cryo-EM structures (Li et al., 2019; Natchiar et al., 2017) and our molecular dynamics (MD) simulations implicate the m_1_acp_3_Ψ-modification in a direct interaction with P-site tRNA. The modification carboxyl-moiety forms a hydrogen bond with the universally conserved RPS16 p.R146 (Jindal et al., 2019), an interaction aided by the m^1^ modification which stabilizes base rotation angle (Fig. S4c). Unlike *TSR3*, the m^1^ transferase of 18S.1248, *EMG1*, is essential and its mutation causes Bowen-Conradi Syndrome (OMIM: 211180, Armistead et al., 2009; Wurm et al., 2010), supporting that 18S.1248 modification plays a crucial role in P-site function.

**Figure 2:**
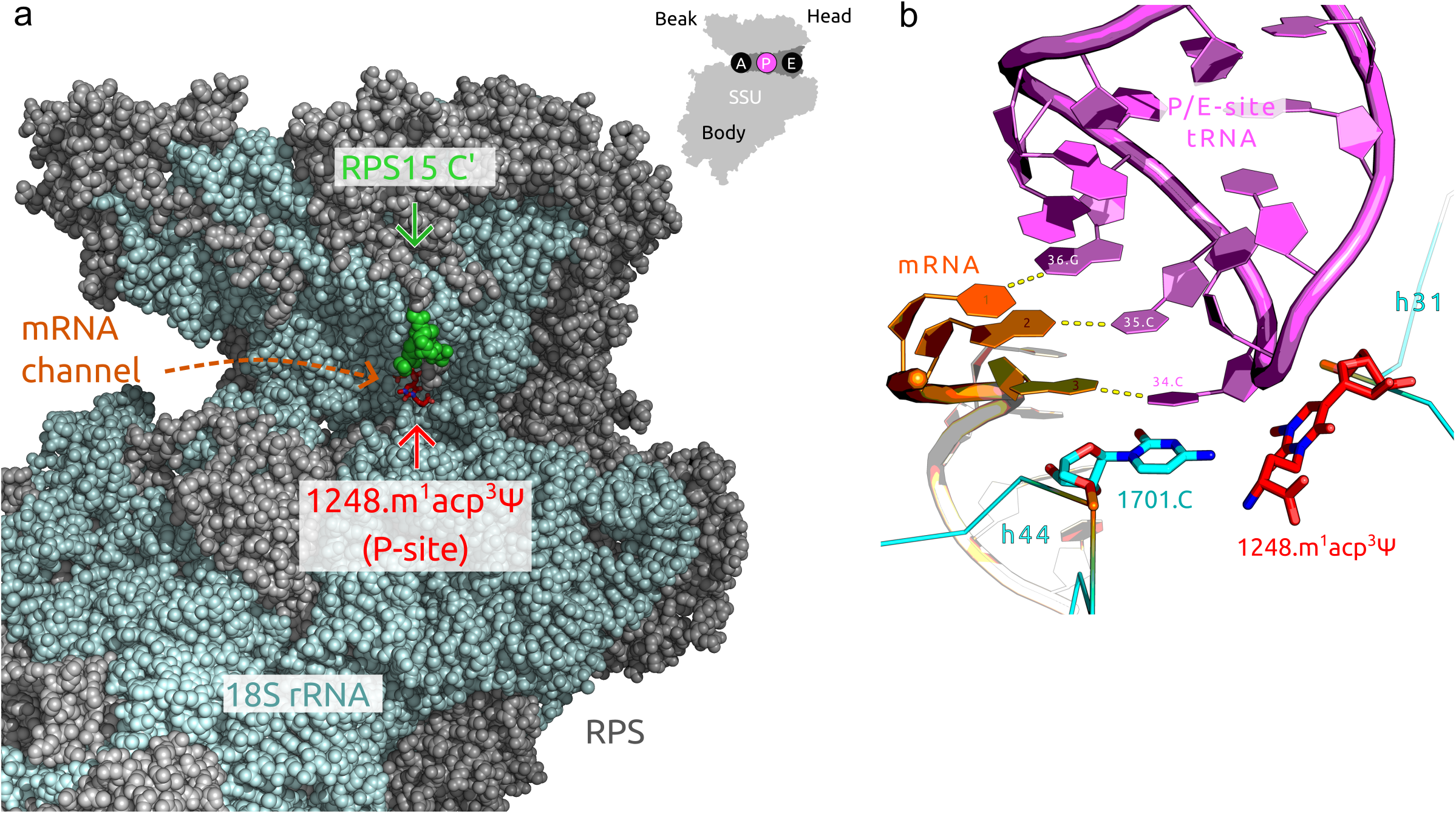
18S:1248.m^1^acp^3^Ψ is located at the peptidyl decoding site. **a**, The mRNA channel of the human small sub-unit (SSU) cryo-EM structure with resolved base modifications (PDB: 6EK0 (Natchiar et al., 2017)). The 18S:1248.m^1^acp^3^Ψ (red) nucleotide is on the loop of the universal helix 31, exposed to the mRNA channel at the center of the P-site. The CLL driver mutations in the RPS15 C’ tail (green) are <12.8Å from 18S:1248.m^1^acp^3^Ψ. The minimal distance is likely shorter as the 10 C’ terminal residues of RPS15 which extend into the P-site are labile and not modeled. **b**, Cryo-EM structure with a P/E-site tRNA, 18S.1248.Ψ and 18S.1701.C base stack with the ribose and base of the tRNA.34.C, respectively (PDB: 6OLE (Li et al., 2019)). 18S.1248.m^1^acp^3^Ψ modification contributes to P-site decoding site stability via interaction with P-site tRNA and RPS16 (Fig. S4).

Start AUG-codon selection and translational initiation is a rate-limiting step in protein-synthesis and both occur at the P-site. Thus, the hypo-m^1^acp^3^Ψ phenotype may demarcate a class of ‘onco-ribosome’ with deregulated translation. It is noteworthy that the two largest effect size RP cancer driver mutations also occur at the ribosomal P-site/tRNA interface, the RPL10 p.R98S at the peptidyl transfer site (Girardi et al., 2018; Kampen et al., 2019) and RPS15 C’ tail mutations adjacent (<12Å) to 1248.m^1^acp^3^Ψ (Fig. 2a) suggesting that the ribosomal P-site is a convergent multi-cancer oncogenic hot-spot.

Since the discovery of streptomycin in 1944, the ribosome has been the target of several important classes of drugs (McCoy et al., 2011). The pervasive and recurrent loss of a solvent-exposed and charged acp_3_ modification at the decoding core of the small sub-unit raises the possibility that this pocket may be therapeutically exploited with ribosome targeting antibiotics or their derivatives as a new generation of chemotherapies.

### Loss of 18S.1248.m^1^acp^3^Ψ modification induces ribosomal protein mRNA translation

To delineate the function of 18S.1248.m^1^acp^3^Ψ, we generated *TSR3* knockout CRC cell lines (HCT116). Similar to yeast (Meyer et al., 2016), *TSR3* is non-essential and we isolated two *TSR3* homozygous knockouts (*TSR3*[KO 1,3]), a heterozygous knockout (*TSR3*[Het 2]), as well as three wildtype control clones (WT[1-3]). Knockouts were functionally confirmed by five independent m^1^acp^3^Ψ assays and editing sites validated by RNA-seq, with *TSR3*[Het 2] showing an intermediate or hypo-m^1^acp^3^Ψ phenotype (Fig. 3a, S2g, S5). The hypo-m^1^acp^3^Ψ seen in *TSR3*[Het 2] is more severe than seen in CRC cell lines (Fig. S2e), but comparable to what is observed in CRC patient tumors (Fig. S1a). SCARLET shows the presence of the Ψ and m^1^Ψ precursor bases at 18S.1248 in *TSR3*[KO 1] cells (Fig. S2h). Together this demonstrates that loss of the acp_3_ modification via *TSR3*[KO] is necessary for nucleotide misincorporation during RT as measured by RNA-seq, supporting hypo-m^1^acp^3^Ψ tumors contain sub-stoichiometric loss of the acp_3_-moiety.

**Figure 3:**
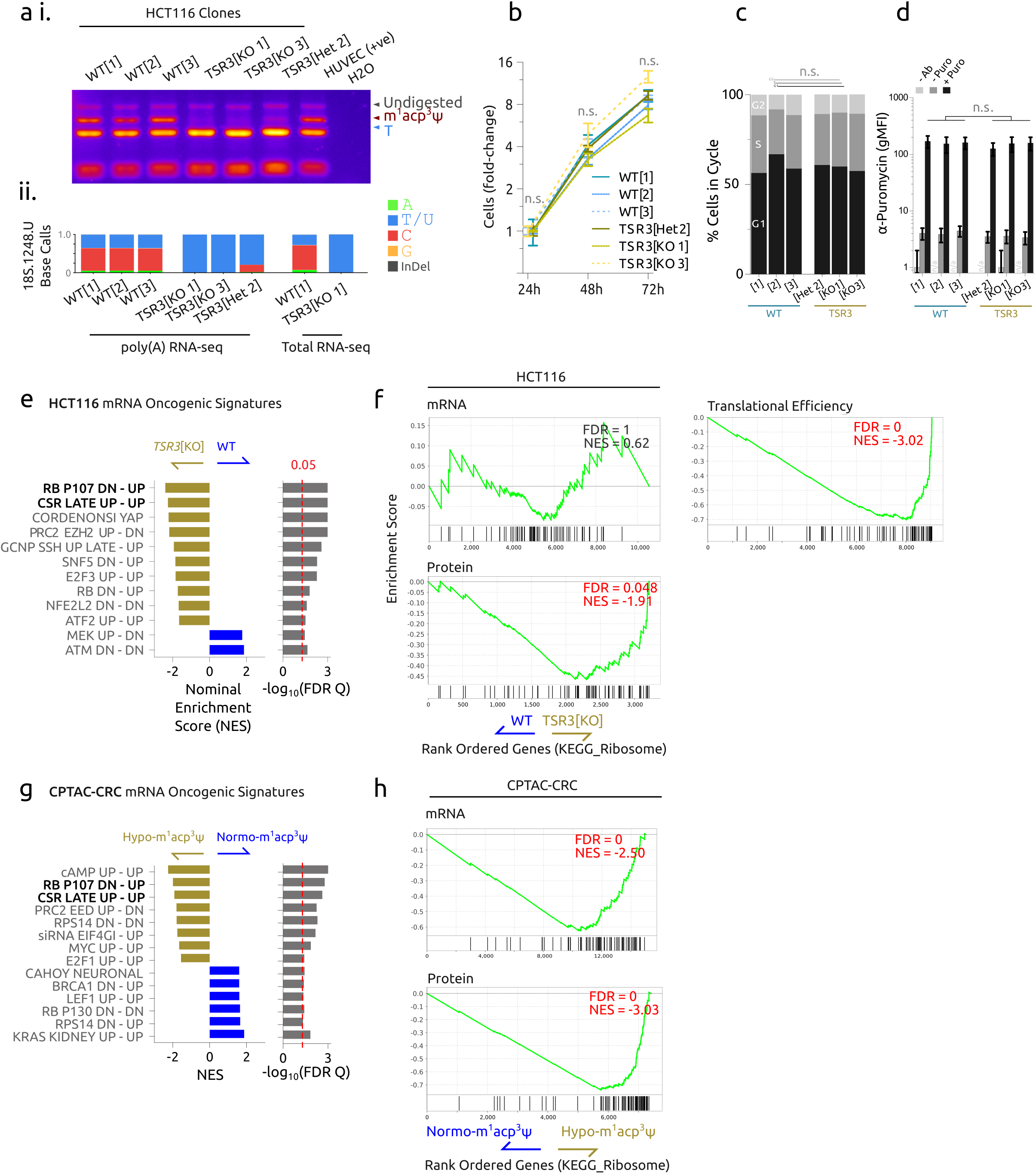
The translational signature of m^1^acp^3^Ψ-deficient ribosomes. **a i**, Reverse transcription (RT)-PCR m^1^acp^3^-assay (see: methods and Fig. S2) and **ii**, RNA-seq measurement for nucleotide misincorporation at 18S:1248.m^1^acp^3^Ψ in clones of the colorectal cancer HCT116 cell line with HUVEC as a normal positive control. **b**, The fold-change growth of HCT116 clone populations in culture, normalized to cells at 24 hours. **c**, Cell cycle timing analysis of log-growth HCT116 cell genotypes. **d**, Total protein translation rates measured by puromycin-incorporation and flow cytometric quantification. **e**, Summary of Gene Set Enrichment Analysis (GSEA) of RNA-seq comparing HCT116 WT[1-3] versus HCT116 *TSR3*[KO 1,3 / Het 2] clones. Only significant (False Discovery Rate Q-value < 0.05) gene sets in the Oncogenic Signature collection are shown. **f**, *TSR3*[KO/Het] cells GSEA shows no change in ribosomal protein (RP) mRNA abundance, while RP translational efficiency, and total protein increases. **g**, Summary of Oncogenic Signature GSEA comparing CPTAC colorectal carcinoma (CRC) tumors with normo-m^1^acp^3^Ψ and hypo-m^1^acp^3^Ψ modification. Gene sets common to HCT116 *TSR3*[KO]/[Het] and hypo-m^1^acp^3^ CPTAC-CRC patients are bolded and **h**, a similar increase in RP abundance is observed.

Morphologically, the HCT116 clones were indistinguishable and showed comparable, rapid growth, cell cycle timing, and loss of TSR3 did not alter global protein translation rates (Fig. 3b-d). To determine how the loss of 18S.1248.m^1^acp^3^Ψ modification alters translation we characterized the transcriptome (RNA-seq) and translatome (ribo-seq) of these cells and validated our findings by global mass spectrometry (Fig. S5-S7).

Transcriptionally, gene set enrichment analysis (GSEA) revealed that *TSR3*[KO]/[Het] (vs. WT) cells were dominated by a proliferative tumor expression signature, characterized by Rb-depletion/elevated-E2F transcription factor activity (Fig. 3e, Fig. S8, table S3). Yet, *TSR3*[KO]/[Het] cells also have a paradoxical decrease in translation of the same E2F target genes (Fig. S8), stressing the specificity of using RNA-expression gene-set classifiers.

To determine how loss of 18S.1248.m^1^acp^3^Ψ modification alters translation, we contrasted translational efficiency between genotypes. *TSR3*[KO]/[Het] cells showed a remarkable enrichment (vs. WT) in the translation of RPs, such as RPS8 and RPL4, with an associated with a depletion of RP mRNA (Fig. 3f, S7d-g, S9), yet overall translation rates and polysome profiles of *TSR3*[KO]/[Het] cells was not altered (Fig. 3d, S9e).

To validate if this RP mRNA/protein signature is present in cancer patients, we analyzed the CPTAC-CRC cohort with tumor matched RNA-seq and proteomics data (Vasaikar et al., 2019). Similar to *TSR3*[KO]/[Het] cell lines, hypo-m^1^acp^3^Ψ CRC tumors share the same E2F oncogenic gene signature and a proteomic increase in RPs relative to normo-m^1^acp^3^Ψ CRC tumor controls (Fig. 3g,h).

There are two hypotheses with which to interpret the hypo-m^1^acp^3^Ψ phenotype. The ‘oncogenic driver hypothesis’ is that m^1^acp^3^Ψ-deficient rRNA arise in tumorigenesis, and their dysregulated translation confers a selective advantage to the cancer, likely via high RP output. The recapitulation of the *TSR3*[KO]/[Het] multi-omic phenotype in hypo-m^1^acp^3^Ψ CRC patients supports a causal model. Alternatively, m^1^acp^3^Ψ-deficiency arises in consequence to hyper-proliferation and high ribosomal biogenesis. Rapid cellular turn-over (oft associated with upregulated E2F1 activity) in turn results in ‘incomplete modification’ of rRNA. Under this model the consequences of m^1^acp^3^Ψ-deficient rRNA is near-neutral or tolerably detrimental to tumor fitness. Nevertheless, hypo-m^1^acp^3^Ψ is a highly recurrent perturbation to the ancient peptidyl-decoding core and underlies a greater cancer-translational phenomenon.

### Loss of m^1^acp^3^Ψ ribosomal RNA modification is a major feature of cancer

Ribosomes are the fulcrum in the central dogma of molecular biology. Multi-omics studies have repeatedly highlighted the discordance between mRNA and protein abundance (Liu et al., 2016; Vogel and Marcotte, 2012), emphasizing the role of translational variability in physio-normal and pathological states. Several recent studies have begun to resolve the ribosome from a uniform assembly into a rich tapestry of functionally heterogeneous complexes making up distinct translational compartments in the cell (2015; Dinman, 2016; Slavov et al., 2015; Xue et al., 2015; Shi et al., 2017; Fujii et al., 2018). Specifically, several rRNA modifications such as pseudouridylation (Ruggero et al., 2003; Jack et al., 2011), ribose 2’-O-methylation (Marcel et al., 2013; Krogh et al., 2016) and, C^5^-methyl cytosine (Heissenberger et al., 2019), have been implicated in altering the translational capacity of cancer cells (reviewed in (Jonkhout et al., 2017; Bastide and David, 2018). Together now with m^1^acp^3^Ψ this supports a broader view of rRNA as a central oncogene, perturbed not genetically, but at the epigenetic modification level. The variable and sub-stochiometric loss of m^1^acp^3^Ψ in cancer is noteworthy and suggests a sub-fraction of a cancer cell’s ribosomes are affected. Further, the loss of 18S.1248 acp^3^-modification altered only a sub-set of (RP) mRNA. Together this suggests that the molecular and possibly tumorigenic function of acp^3^-modification is mediated by a distinct class of ribosome, although further research is needed to resolve this mechanism.

We have discovered a pervasive and cancer-specific ‘onco-ribosome’ marked by loss of rRNA m^1^acp^3^Ψ modification. Enticingly, the cancer-specific m^1^acp^3^Ψ-deficient ribosomes are exceptionally recurrent and can be explored as a novel chemotherapeutic class.

## Data and code availability

Sequencing data generated in this study is available on NCBI Short Read Archive (study: PRJNA602544). Proteomics data generated in this studyare available on PRIDE (accession pending). Electronic laboratory notebook for these experiments and analysis scripts are available at https://www.github.com/ababaian/Crown.

## Acknowledgments

AB and KR thank Ada Maple Babaian for helpful discussion on the manuscript. This work was supported by grants from the Natural Sciences and Engineering Research Council of Canada (NSERC) and the Leukemia and Lymphoma Society of Canada to DLM. AB was supported by a National Science and Engineering Research Council (NSERC) Alexander Graham Bell Graduate Scholarship and a Roman Babicki Fellowship in Medical Research from the University of British Columbia. Sequence computation was provided by Amazon Web Services (AWS) under a Research Grant to AB. CRISPR-Cas9 reagents were provided by the Integrated DNA Technologies under the CRISPR-Challenge Prize to AB/DLM. Molecular dynamic computation was provided by Compute-Canada. DG was supported by an Alberta Innovates (Technology Futures) Graduate Student Scholarship. HJW was supported by an Alberta Innovates (Strategic Chairs Program SC60-T2) and NSERC Discovery Grant (RGPIN-2016-05199).

Data used in this publication were generated by The Cancer Genome Atlas (TCGA) Research Network (https://www.cancer.gov/tcga), National Cancer Institute Clinical Proteomic Tumor Analysis Consortium (CPTAC) and Genentech/gRED.

## Author contributions

A.B. discovered hypo-m^1^acp^3^Ψ led the study and. A.B. performed DNA- and RNA-seq analysis. A.B. and K.R. performed molecular and cell culture experiments. A.B. invented the aRT-PCR assay for m^1^acp^3^Ψ. S.D. performed primer extension and aRT-PCR optimization. D.G. performed molecular dynamic simulations and H.J.W. and A.B. helped analyze the data. I.M. prepared ribo-seq libraries, performed SCARLET validation and invented the aminocarboxylpropyl biotinylation method. A.B. and M.M processed and analyzed the ribo-seq data. S.E.S.M. performed the LC-MS/MS. H.J.W., M.L., G.M., and D.L.M. provided expert advice for experiments. All authors contributed to the design of the study. A.B. prepared the manuscript and figures.

## Correspondence and requests for materials

Should be addressed to A.B.

## Supplementary Information

Is available for this paper

---

**Supplementary Figure 1: Detailed look at hypo-m**^**1**^**acp**^**3**^**Ψ in the TCGA cohorts**

**a**, The 18S.1248.U variant allele frequency (VAF) from 33 TCGA patient cohorts (study abbreviations in table S1). **b**, The rDNA position underlying *18S.*1248.U (*RNA45S.*4908.T) is invariable in TCGA-COAD. Matched tumor-normal whole exome sequencing (WXS) data from 438 TCGA-COAD patients was aligned to *hgr1.* 415 cancer and 413 normal samples contained rDNA coverage of *RNA45S.*4908.T. Three normal samples had a single variant read each at this position, consistent with sequencing error. **c**, Batch-specific shift in the average 18S.1248.U VAF in i, TCGA-COAD and ii, TCGA-DLBC libraries. Similar to seen in the aRT-PCR assay for m^1^acp^3^Ψ (Figure S2), there are batch-effects with m^1^acp^3^Ψ misincorporation, but the relative decrease in m^1^acp^3^Ψ-modification in CRC compared to normals is seen in across all batches. **d**, The gene expression of the m^1^acp^3^Ψ modifying enzymes *TSR3* and *EMG1* is not decreased or lost across the TCGA cohorts.

**Supplementary Figure 2: Aminocarboxyl propyl Reverse Transcription (aRT)-PCR assay for m**^**1**^**acp**^**3**^**Ψ and rRNA modification in cell lines**

**a i**, The 18S.1248.m^1^acp^3^Ψ modification assay is based on the misincorporation of nucleotides in first strand complementary DNA (cDNA) strand synthesis by reverse transcriptase (RT). The cDNA is then PCR amplified and **ii**, the ratio of reference T and not-T (V = A, C, or G) is genotyped by the *HinFI* restriction enzyme cut-site which overlaps 18S.1248. **b**, The choice of RT-enzyme; SuperScript III (SSIII), SuperScript IV (SSIV), WarmStart RTx (WS RTx) or, UltraScript 2.0 (US 2.0), influences nucleotide misincorporation rates and the variant allele frequency (VAF) read-out of the m^1^acp^3^-assay, although VAF remains consistent across biological replicates of input RNA of the colorectal cancer (CRC) cell line HCT116 wildtype clone 1 (WT[1]), or HCT116 with *TSR3* gene knockout clone 1 (*TSR3*[KO 1]). **c**, PCR replicates of WT[1] and TSR3[KO 1] cDNA, shows consistent readout. **d**, HCT116 WT[1] and *TSR3*[KO 1] RNA was mixed at fixed weight ratios (μg total RNA) prior to RT to determine if the assay is quantitative for m_1_acp_3_ modification. **e**, The aRT-PCR assay applied to 11 CRC cell lines, primary human umbilical vein endothelial cells (HUVEC) as a normal control and, the blast-phase chronic myelogeous leukemia cell line K562 as a hypo-m^1^acp^3^Ψ positive control. **f i**, Polysomal fractionation and **ii**, sub-fraction m^1^acp^3^Ψ RT-PCR assay of the hypo-m^1^acp^3^Ψ cell line K562. In cells containing a mixture of +/- m^1^acp^3^Ψ modification, unmodified rRNA incorporates into mature ribosomes and is enriched in the lower-order mono- and di-somes. **g**, Primer extension assay for 18S.1248.m^1^acp^3^Ψ modification in HCT116 WT[1] and *TSR3*[KO 1]. The helix 31 structural stop and rRNA truncation via DNA cut oligo + RNase H treatment is used as internal load controls. **h**, SCARLET assay of HCT116 WT[1] and *TSR3*[KO 1] is not suitable for measuring m^1^acp^3^Ψ as this base may interfere with probe hybridization, but precursor bases were detected and accumulated in the TSR3 knock-out cells. **i**, Total RNA treated with the amino-reactive ester, N-hydroxysuccinimidobiotin and detected with **i.** ethidium bromide staining as load control and **i.** anti-streptavidin-HRP antibody and. *TSR3*[KO 1] cells show loss of primary amine reactivity (such as m^1^<u>a</u>cp^3^Ψ) in 18S rRNA but not in tRNA.

**Supplementary Figure 3: 18S.1248.m**^**1**^**acp**^**3**^**Ψ is absolutely conserved in *Eukarya***

**a**, Evolutionary conservation of 18S rRNA in the locus surrounding helix 31 in *Eukarya* and select *Archaea* and *Eubacteria* species (Bernier et al., 2014; Petrov et al., 2014). Magenta arrow indicates the position homologous to human18S:1248.U. **b-d**, The conserved secondary structure of helix 31 and its known modification sites (Machnicka et al., 2013; Petrov et al., 2014).

**Supplementary Figure 4: Ribosomal molecular dynamics and modification modeling**

**a**, The m^1^acp^3^Ψ modification stabilizes the decoding peptidyl (P-) site via a hydrogen bond with the universally conserved RPS16 (uS9) p.R146 residue. Whole ribosome molecular dynamics simulations (MD) were ran for up to 50ns with 18S.1248.m^1^acp^3^Ψ, acp^3^Ψ, m^1^Ψ, Ψ or U base in an empty (**b**,**d**, PDB: 6EK0) or tRNA-occupied P-site (**c**,**e**, PDB: 6OLE). **b**,**i.** Root mean squared deviation (RMSD) of empty P-site MD atoms indicate simulations stabilize after 5ns (with up to ∼6Å RMSD), up to 45ns (yellow highlight) was used for analysis. **ii**, Minimal distance between m^1^acp^3^Ψ (3-carboxyl oxygen) or uridine (4-oxygen) and the closest guanidinium hydrogen of RPS16 p.R146 supports that m^1^ and acp^3^ are necessary to be within 3.5Å for hydrogen bonding and nucleotide stabilization. **c**, RMSD of P-site occupied simulations with **ii.** Root mean squared fluctuation (RMSF) of 18S rRNA adjacent to 18S.1248 and **iii**, within helix 31 shows 18S.1248.m^1^acp^3^Ψ is comparable relative to other modification states. The average (**iv**) and standard deviation of **χ**-angle of the 18S.1248 nucleotide in each simulation shows acp^3^ increases torsional variation of the base which is stabilized by the m^1^ modification. **d**, 20ns simulation showing bonding between 18S.1248.m^1^acp^3^Ψ and RPS16 p.R146 compared to **e**, the structure with mRNA and P-site tRNA. We postulate that 18S.1248 acp^3^-modification is involved in coordinating RPS16 p.146 for P-site tRNA positioning and contributes to the stability of the decoding core. Statistical difference between **χ**-angle mean (Tukey HSD), and standard deviation (Benjamini-Hochberg corrected F-tests) were used (* is p < 0.05, ** is p < 0.001 and *** is p < 0.0001).

**Supplementary Figure 5: RNA-seq and *TSR3* editing site validation**

HCT116 WT[1-3] versus *TSR3*[KO]/[Het] RNA-seq metrics. **a**, Differential mRNA expression of expressed (reads per million kilobase, RPKM_mRNA_ >0.1) between WT[1-3] and *TSR3*[KO]/[Het] clones. **b**, Hierarchical clustering of libraries based on expressed genes. Globally, *TSR3*[Het 2] is more dissimilar to *TSR3*[KO 1,2] clones. **c**, MA-plot for mRNA expression highlighting genes in the ‘*KEGG_RIBOSOME’* and ‘*RB_P107_DN.V1_UP*’ gene sets (see: Fig 3). **d**, Volcano plot for differentially expressed genes between WT and *TSR3*[KO]/[Het] with statistically significant genes highlighted in red (list is available in Table S5). **e**, *TSR3*[KO]/[Het] lines have decreased *TSR3* expression measured by RNA-seq (in reads per kilobase per million mapped reads, RPKM). **f**, *TSR3* gene structure and IGV screenshots showing the editing sites in (**g**) *TSR3*[KO 1,3], and (**h)** *TSR3*[Het 2].

**Supplementary Figure 6: Ribo-seq**

HCT116 WT[1-3] versus *TSR3*[KO]/[Het] ribo-seq metrics. **a, i**, As a quality control metric, the length distribution of mapped ribosome-protected fragments for each of the two WT[1-3], and three *TSR3*[KO]/[Het] biological replicates of ribo-seq libraries were plotted. The *TSR3*[KO 3] biological replicate 2 (r2) library had a bi-modal read-length distribution, peaking at 22 and 28 nt suggesting incomplete cycloheximide treatment (Lareau et al., 2014) thus, this library was excluded from downstream expression and positional analyses. **ii**, Short (21-23 nt) ribosome fragments coincide with ribosomes stalled in the rotated, post peptide-bond state (Lareau et al., 2014). *TSR3*[KO 1,3] libraries had fewer short fragments implying m^1^acp^3^Ψ-deficient ribosomes have a lower probability of being in the rotated transition state relative to WT ribosomes. **b**, Hierarchical clustering of libraries based on total translation recapitulates mRNA clustering. **c**, Metagene heatmap of P-site occupancy around start and stop codons. For each read optimal P-site offset was determined (13 nt from 5’ end) and P-site occupancy computed. Log10-transformed P-site coverage is shown for all replicates, scaled by min and max number of reads per sample. A clear trinucleotide periodicity in the CDS of transcripts is observed. **d**, Differential translation of ribo-seq expressed (RPKM_Ribo_ > 0.1) genes. **e**, MA-plot and **f**, volcano-plot for total translation, highlighting the ribosome and RB/E2F gene sets. **g**, The log_2_ mRNA fold-change and log_2_ translation fold-change (WT / *TSR3*[KO]/[Het]) and detail inlay, with each gene size-scaled by mean RPKM_mRNA_. RB/E2F gene sets are highlighted, showing RP genes are more efficiently translated in *TSR3*[KO]/[Het] clones (all points below diagonal in, see also Fig 3). **h**, The P-site periodicity within each library showed the majority of CDS ribosomes were in-frame, with no significant difference in frame-shifting upon m^1^acp^3^Ψ perturbation. **i**, P-site occupancy was calculated over all expressed coding sequences (CDS). Globally, there was no significant difference between P-site occupancy per codon in WT[1-3] in *TSR3*[KO]/[Het] libraries. Since 18S.1248.m^1^acp^3^Ψ is located at the P-site where initiation codon selection occurs, we tested if the initiation AUG codon was differentially occupied between any genotypes. *TSR3*[Het 2] and *TSR3*[KO 1], but not *TSR3*[KO 3] have elevated AUG occupancy relative to WT clones supporting a slower global initiation rate in those two samples. Tukey HSD test was used for testing a statistical difference between group means (* is p < 0.05, ** is p < 0.001 and *** is p < 0.0001).

**Supplementary Figure 7: Liquid-Chromatography Tandem Mass Spetrometry (LC-MS/MS) validation**

HCT116 WT[1-3] versus *TSR3*[KO]/[Het] LC-MS/MS results. **a**, After multiple-testing correction, no proteins were significantly different between genotypes, and no TSR3 peptides were found in the samples. **b-d**, Volcano plots for RNA-seq, ribo-seq, translational efficiency (Ribo-seq_RPKM_/RNA-seq_RPKM_) and proteomic data with genes in the ‘KEGG_RIBOSOME’ gene-set highlighted. As a complimentary analysis to GSEA, a Fisher’s Exact Test was performed to test for enrichment of ‘KEGG_RIBOSOME’ genes either up- or down-regulated between genotypes, compared to all other genes.

**Supplementary Figure 8: The RB/E2F transcriptional signature associated with *TSR*[KO]/[Het]**

HCT116 WT[1-3] versus *TSR3*[KO]/[Het] Gene Set Enrichment analysis (GSEA) for the E2F Targets and oncogenic signature of *RB1, RBL1* (p107) knock-out. **a**, Rb proteins are repressors of the E2F transcription factors. The of E2F GSEA (**i**) was highly enriched upon *TSR3*[KO]/[Het] with (**ii**) *TSR3*[Het 2] showing the strongest enrichment. **b**, Gene set mRNA expression is enriched specifically in genes upregulated upon Rb1 and Rb1;p107 knockout (gene sets: *RB_DN.V1_UP, RB_P107_DN.V1_UP*). **c**, The translational output (ribo-seq signal) of genes upregulated in Rb1;p107 knockout remains increased but **d**, this gene set is translated less efficiently and, **e**, has no difference at the proteome level.

**Supplementary Figure 9: Representative changes to translational efficiency**

Ribo-seq traces for representative genes **a**, *EEF1A2* (eukaryotic translation elongation factor 1 alpha 2), **b**, *RPS8* (ribosomal protein S8), and **c**, *RPL4* (ribosomal protein L4) for HCT116 WT[1-3] versus *TSR3*[KO]/[Het]. **d**, Matched RNA expression (in reads per kilobase per million mapped reads, RPKM), Ribo-seq CDS expression (RPKM), translational efficiency (Ribo-seq_RPKM_/RNA-seq_RPKM_), and proteomic expression (median normalized expression values). **f**, Polysome profiles and relative area under the curve (AUC) intensity for sub-fractions (highlighted in gray) (inlay). Total absorbance was normalized to 100 artificial units in the working range of the gradient (between red ticks on x-axis).

**Supplementary Table 1:**

Accessions of DNA and RNA sequencing libraries used in this study. 18S.1248.U variant allele frequency (VAF) is provided for each sample.

**Supplementary Table 2:**

Primers, oligos, and guide RNA sequences.

**Supplementary Table 3:**

Significant (FDR q-value < 0.05) Gene Set Enrichment Analysis results for differential transcriptomics in HCT116 WT[1-3] versus *TSR3*[KO]/[Het] with the Hallmarks, C3 promoter motif, and C6 oncogenic signatures gene sets.

**Supplementary Table 4:**

Significant (FDR q-value < 0.05) Gene Set Enrichment Analysis results with the C2 gene-ontology and pathways gene sets for differential transcriptomics and translational efficiency in HCT116 WT[1-3] versus *TSR3*[KO]/[Het] cell line and transcriptomics and proteomics in normo-m^1^acp^3^Ψ versus hypo-m^1^acp^3^Ψ CPTAC-CRC tumors.

**Supplementary Table 5:**

Significant (adjust P-value < 0.05) genes deferentially expressed between WT[1-3], and *TSR3*[KO 1,3]/ [Het 2] in the RNA-seq and Ribo-seq experiments.

## Extended Methods

### Ribosomal sequence alignment and variant allele frequency calculations

RNA-seq libraries used in this study were prepared via poly-A selection to enrich for the ∼5% of mRNA from total RNA. Since rRNA is ∼80% of cellular RNA it invariably ‘contaminates’ RNA-seq libraries. Typically, poly-(A) RNA-seq libraries contain 3.55% (+/- 0.685, 95% CI, CRC I cohort, N = 66) of total reads aligned to rDNA. Whole genome DNA-seq (WGS) libraries were processed from initial fastq files. The 876 TCGA-COAD whole exome DNA-seq (WXS) aligned libraries were downloaded and reads mapping to the contig “chrUn_GL000220v1” (which contains a complete *RNA45S* gene) were extracted with ‘*samtools*’ into fastq files. 415/438 cancer samples and 413/438 normal samples had rDNA coverage at a total of 15,175x and 17,936x, respectively. A complete list of library accessions used in this study is available in Table S1 (Cancer Genome Atlas Research Network et al., 2013; Ghandi et al., 2019; Lim et al., 2015; Seshagiri et al., 2012).

Libraries were aligned to the *hgr1* reference rDNA sequence (Babaian, 2017) with *bowtie2* (v. 2.3.5.1, command: *‘bowtie2 --very-sensitive-local -x hgr1 -1* <*read1.fq.gz*> *-2* <*read2.fq.gz*>*’*) (Langmead and Salzberg, 2012). For each cohort of libraries, a genomic variant call format (GVCF) was created with *bcftools* (v. 1.9, command: ‘*bcftools mpileup -f hgr1.fa –max-depth 10000 -A -min-BQ 30 –b* <*bam.file.list*>*’*) (Danecek et al., 2011).

GVCF files were processed in R by custom scripts to calculate variant allele frequency (VAF). VAF is defined as 1 – reference allele frequency (reference allele depth of coverage / total depth of coverage) (scripts available at https://www.github.com/ababaian/crown)

The threshold to define hypo-modification of an RNA base (including 18S.1248.m^1^acp^3^Ψ) was defined as three standard deviations below average VAF of the normal samples within the same cohort (false discovery rate = 0.00135) when available. Fixed formalin paraffin embedded (FFPE) libraries in TCGA were negative for 18S.1248.m^1^acp^3^Ψ, 28S.1321.m^1^A and 28S.4532.m^3^U modification signatures and excluded from further analysis. In the CPTAC-CRC cohort (normal RNA-seq is unavailable), hypo-m^1^acp^3^Ψ and normo-m^1^acp^3^Ψ was defined by the lower (<25%) and upper (>75%) quantiles of samples within a batch.

### Transcriptome and translatome alignment, assembly and differential expression

RNA-seq reads were aligned to *hg38* (*GRCh38)* reference genome with *tophat2* (v.2.0.14) (Kim et al., 2013). Individual transcriptome assemblies for HCT116 [WT 1-3], [KO 1,2] and [Het 2] libraries were generated with *stringtie* (v 2.0) (Pertea et al., 2015), and then all merged together with the human *gencode basic* gene annotation (v. 31) (Frankish et al., 2019) ultimately yielding the *hct116_gencode.v31* reference gene set.

To generate a single-copy reference transcriptome for ribo-seq analysis of HCT116, isoform-specific quantification of gene expression was performed on the *hct116_gencode.v31* gene set with ‘*stringtie -G hct116_gencode.gtf’*. For each gene with non-zero expression (>10 unique reads), the one highest expression isoform (average expression from each clone) was chosen as the reference transcript for that gene.

For ribo-seq alignment, after read adapter trimming and alignment to *hgr1* as above, unmapped reads were aligned against a containment file containing human tRNA, mtDNA, snoRNA, snRNA and miRNA sequences. Reads remaining unmapped were then aligned to *hg38* and the *hct116_transcriptome* with *STAR* aligner (v. 2.5.2b, command: *‘STAR --genomeDir hg38 --readFilesIn*

<*input.fq*> *--sjdbFileChrStartEnd hg38/sjdbList.out.tab --outFilterMultimapNmax 10 -- outFilterMismatchNmax 5 --outFilterMatchNmin 15 --alignSJoverhangMin 5 –seedSearchStartLmax 20 --outSJfilterOverhangMin 30 8 8 8 --quantMode TranscriptomeSAM’*) (Dobin et al., 2013). Transcriptome aligned Ribo-seq data was analyzed in *R* (v. 3.5.1) using the *riboWaltz* package (v. 1.1.0) (Lauria et al., 2018).

Gene-level expression and total translation was quantified with the *DEseq2* (Love et al., 2014) R package using *hg38* aligned bam files and the *hct116_gencode.v31* reference gene set. Ribo-seq Gene expression (RPKM) was calculated based only on reads mapping in-frame to a gene CDS, measure translating ribosomes. Translational efficiency was calculated per genotype as log2(Ribo-seq Gene_RPKM_ / RNA-seq Gene_RPKM_).

Gene expression and translation differences were calculated by Gene Set Enrichment Analysis (*GSEA*, v.4.0.0) (Subramanian et al., 2005) with ‘*-permute gene_set -nperm 5000’* and standard parameters. Transcriptomic GSEA was performed using the MSigDB (Liberzon et al., 2015) (v 7.0): hallmark, C2 pathways, C3 motif search, and C6 oncogenic signatures gene sets. Translatomic and proteomic GSEA was performed with C5 Gene Ontology (GO) gene set.

All bioinformatic analyses were scripted for reproducibility and are available at https://www.github.com/ababaian/crown, RNA-seq and Ribo-seq data can be directly viewed in the UCSC genome browser at https://genome.ucsc.edu/s/Artem%20Babaian/HCT116_TSR3%2DKO.

### HCT116 cell culture and TSR3 knockout

The colorectal carcinoma cell line HCT116 (CCL-247, ATCC, Manassas, VA) was cultured in complete media [DMEM media (#36250, STEMCELL Technologies, Vancouver, Canada) supplemented with 10% fetal bovine serum (F1051, Invitrogen, Waltham, MA)].

To generate *TSR3* knockouts, 10^5^ HCT116 cells were transfected with 10 nmol of one of three *TSR3* targeting Alt-R CRISPR-Cas9 ribonucleoproteins or non-targeting controls (table S2) by manufacturer’s protocol (1081059, Integrated DNA Technologies (IDT), Coralville, IA). After 24 hours, single cells from each treatment group were isolated by limiting dilution and confirmed to be 1 cell /well by microscopy. Single cell clones were expanded to 5×10^5^ cells at which point half the culture was frozen (culture media + 10% DMSO) and half were processed for RNA. *TSR3* knockouts were genotyped by RNA-seq and functional knockout was confirmed by three independent m^1^acp^3^Ψ assays (figure 3, S2).

Cell lines and clonal isolates were tested to be free of mycoplasma contamination by DAPI staining and microscopy and with LookOut Mycoplasma Detection Kit (MP0035, MilliporeSigma, Burlington, MA) by manufacturer’s protocol.

### Cell cycle and puromycin incorporation assays

HCT116 cells were seeded at 2×10^5^ cells per 6-well plate and allowed to grow until log-phase of growth at ∼50-75% confluency. To detach cells, cells were washed twice with PBS, detached with 0.5 ml 0.25% trypsin EDTA (07901, STEMCELL) for 5 minutes at RT, followed by inactivation with 2.5 ml of complete media and cells pelleted at 300 x g for 5 minutes at 4°C. Pellets were washed twice with cold PBS.

For cell cycle analysis, cells were fixed overnight at 4°C in 70% EtOH solution. Fixed cells were washed in PBS and resuspended in 500 ul of staining solution [Propidium Iodide (50 ug/ml), RNAse A (0.1 mg/ml), and 0.05% Triton X-100 in PBS], and incubated 30 minutes at RT in the dark followed by a wash step. Flow cytometry was performed on a FACScalibur with *CellQuest* acquisition software (BD Biosciences) acquiring at least 10,000 cells per sample and analyzed with *FlowJo* (v 10.0.8). Cell cycle gating was performed on single-cells using the automated ‘Watson Model’.

Puromycin-incorporation assay was performed as previously described (Schmidt et al., 2009). Briefly, cells were pulsed with either complete media only or complete media containing 10 ug/mL Puromycin (73342, STEMCELL) for 10 minutes in cell incubator. Cells were washed twice in pre-warmed PBS and chased for 50 minutes in incubator. Cells were then detached, washed and fixed in 2% paraformaldehyde in PBS for 15 minutes at RT. Cells were then washed twice in wash solution (5% FBS, and 0.1% Saponin in PBS). 1×10^5^ cells were stained in 100 ul of 1:100 anti-puromycin antibody (12D10) conjugated to Alexa Flour 488 (MAEBE343-AF488, MilliporeSigma) in wash solution for 30 minutes on ice followed by two final washes in wash solution and resuspension in 100 ul PBS for flow cytometry to acquire at least 5,000 cells per sample. Unstained controls were performed on puromycin treated cells. Puromycin staining intensity correlated with natural variation in cell size (forward scatter signal, FSC), thus the 488nm staining signal intensity (anti-puromycin) was normalized by cell size (normalized fluorescence = 250 * anti-puromycin / FSC). The geometric mean florescent intensity (gMFI) and standard deviation for each sample is reported.

### RNA isolation

Cells for RNA extraction were lysed directly in TRIzol reagent (15596-018, Invitrogen), spun 5 min at 12,000 x g to pellet fat and nuclear DNA and then frozen at -80°C. RNA extraction was carried out by manufacturer’s protocol. RNA quality was assessed via 2% denaturing RNA agarose gel electrophoresis (heat treated, 95°C for 5 minutes in 1.5x formamide loading buffer (Masek et al., 2005)) and concentration/purity assessed by spectrophotometer (NanoDrop 2000, ThermoFisher, Waltham, MA). RNA quality for RNA-seq library preparation had a >9.9 RIN score measured by Bioanalyzer 2100 (Agilent, Santa Clara, CA).

### Primer extension for 18S.1248.m^1^acp^3^Ψ modification

Primer extension was performed with 1 µg of total RNA, incubated with 2 pmol of *PE_1248_BLOCK* (IDT) primer and 2U of RNase H (18021-014, Invitrogen) or mock enzyme treatment at 37°C for 20 min followed by heat inactivation at 65°C for 10 min. SuperScript III reverse transcriptase (18080044, lot #2042663, Invitrogen) and the fluorophore labeled *PE_1248_FAM* primer were added for primer annealing and RT (1h at 50°C) as described by Schuster and Bertram (Schuster and Bertram, 2014). Labeled cDNAs were re-suspended in 1.5x formamide loading buffer and heated to 95°C for 3 min to eliminate secondary structures (Masek et al., 2005). Samples were separated on a 2% agarose gel at 114 V for 3h at 4°C or on a 12.5% polyacrylamide gel at 45 mA for 2.5h in 1x TBE. After migration, the gel was visualized with the Typhoon FLA 9500 laser scanner (FAM filter, 50 µm pixel and 450 V unless otherwise noted, GE Healthcare, Chicago, IL).

### Aminocarboxyl propyl Reverse Transcription (aRT)-PCR for 18S.1248.m^1^acp^3^Ψ modification

The aminocarboxylpropyl reverse transcription (aRT)-PCR assay for 1248.m^1^acp^3^Ψ modification was performed with 1 ug of DNase treated (AM1907, lot #00733051, Invitrogen) RNA after total RNA quality was assessed by denaturing agarose gel electrophoresis (Masek et al., 2005). RT reaction was carried out with SuperScript III (Invitrogen), SuperScript IV (18090010, lot #00721480, Invitrogen), UltraScript 2.0 (PB30.31-10, lot #PB130614-01-5, PCR Biosystems, Wayne PA) and WarmStart RTx (M0380L, lot #0061705, NEB) by each manufacturer’s protocol with minor modifications. aRT reactions were carried out with a random hexamer primer only, and not poly(T) oligos. cDNAs were diluted five-fold and used as template for PCR (30 cycles: 94°C for 30 s, 55°C for 30 s, 72°C for 30 s) with *macp_F1* and *macp_R1* primers (table S2). Amplicons were digested with HinFI (R0155S, New England BioLabs) (25°C for 5 sec, 37°C for 90 min, 80°C for 20 min). Samples were separated on a 2.25% agarose gel in 1x TBE at 200 V for 45 min at 4°C. After migration, the gel was post-stained in 1x GelRed (41003, Biotium, Fremont CA) for 30 min. Gels were visualized by UV transillumination, captured in gray scale with a digital camera and pseudo-colored in ImageJ (Schneider et al., 2012) (v 1.52h, Lookup table > Fire) which retains the original pixel intensity values but highlights band-intensity visualization.

### SCARLET for 18S.1248.m^1^acp^3^Ψ modification

The validation of 18S.1248.m1acp3Ψ was performed by site-specific cleavage and radioactive-labeling followed by ligation-assisted extraction and thin-layer chromatography (SCARLET) as previously described (Liu et al., 2013) with minor modifications.

Briefly, 1 μg of total RNA from HCT116 WT or *TSR3*[KO] was annealed with 3 pmol of chimera oligo (oligo sequences in Table S2) in 2.5 μl 30 mM Tris-HCl pH 7.5, by heating at 95 °C for 1 min. The sample was incubated at 44 °C for 1 h in total volume of 5 μl 0.4× T4 PNK buffer (B0201S, New England Biolabs) supplemented with 1 U/μl RNase H (Epicentre, Madison, WI) and 1 U/μl FastAP (EF0651, Thermo Fisher Scientific), followed by incubation at 75 °C for 5 min. 1 μl of radioactive labeling buffer (1× T4 PNK buffer, 2.5 U/μl T4 PNK, 4 μCi/μl [γ-^32^P]ATP) was added to the RNA and incubated at 37 °C for 1 h and then at 75 °C for 5 min. The mixture was then annealed with 4 pmol splint oligo and 4 pmol 116-mer DNA oligo by heating at 75 °C for 3 min followed by addition of 2.9 μl ligation buffer (1.4× PNK buffer, 0.2 mM ATP, 57% DMSO, 5 U/μl T4 DNA ligase) and incubation for 3.5 h at 37 °C. The ligation was stopped by adding an equal volume of RNA urea loading buffer (8 M urea, 100 mM EDTA, 0.025% (w/v) bromphenol blue) and then was digested by incubated with 1 μl RNase T1/A mixture (EN0551, Thermo Fisher Scientific) at 37 °C for 16 h. The sample was loaded on 10% denaturing urea-PAGE to isolate the ligation product, which was desalted by ethanol precipitation. The pellet was resuspended in 9 μl sodium acetate/acetic acid buffer pH 4.6, supplemented with 2mM ZnCl2 and 5 units of nuclease P1, reaction was incubated) at 37 °C for 4 h and then was spotted on a TLC plate for separation. The result was visualized on a Typhoon FLA 9500 phosphor imager (GE Healthcare).

### Biotinylation of 18S.1248.m^1^acp^3^Ψ modification

We have exploited reactivity of the m^1^acp^3^Ψ base with N-hydroxysuccinimide esters which was initially described in early 1970’s (Gillam et al., 1968; Schofield et al., 1970; Friedman, 1972). Briefly, 10 µg of total RNA from HCT116 WT or *TSR3*[KO] in 47 µl of reaction buffer (100 mM phosphate buffer pH 8.0, 150 mM NaCl, 0.5 M EDTA pH 8.0) were denatured for 5 min at 70 °C and placed on ice. After that 1µl of 100mM NHS-dPEG12-biotin (QBD10198-50MG, Sigma Aldrich) in DMSO was added. Reaction was incubated on ice for 2 h, every 40 minutes 1 µl of 100 mM of NHS-dPEG12-biotin was added (3 times in total). After treatment RNA was purified with Zymo-Spin IIC column, resolved on denaturing agarose gel and blotted on Hyperbond N+ (Amersham) membrane. RNA was UV crosslinked twice with 1200 µJ followed by incubation in blocking solution (phosphate buffered saline [PBS], pH 7.5, 10% SDS, 1 mM EDTA) for 20 min. The membrane was probed with 1:10000 dilution of 1 mg/ml streptavidin-horseradish peroxidase (Pierce) in blocking solution for 15 min, washed six times in PBS containing decreasing concentrations of SDS (10%, 1%, and 0.1% SDS, applied twice each) for 5 min and finally the biotin signal was visualized by ECL detection reagent (10 min exposure time).

### RNA-seq

RNA-seq library preparation and sequencing was performed by the BC Cancer Genome Sciences Centre, Vancouver, Canada. Briefly, 75-bp stranded and paired-end poly-(A) RNA-seq libraries were prepared with NEBNext poly(A) mRNA magnetic isolation module (E7490L, New England BioLabs (NEB), Ipswich, MA), Maxima H minus First Strand cDNA synthesis kit (K1652, Thermo-Fisher), and NEBNext Ultra II directional RNA second strand synthesis (E7771, NEB). The total RNA-seq libraries were prepared in parallel but without poly-(A) selection and only 2x PCR cycles (for adapter ligation). Libraries were sequenced on a HiSeq 2500 (Illumina, San Diego, CA).

### Ribo-seq

Ribosome foot printing (ribo-seq) was performed as previously described (Ingolia et al., 2012) with minor modifications. For cell harvesting, the culture medium was aspirated, cells were washed twice with ice-cold PBS supplied with 100 μg/ml cycloheximide and plates were flash-frozen in liquid nitrogen. For cell lysis, the plates were placed on wet ice and 400 µl of mammalian polysome buffer (MPB) [20 mM Tris-HCl, pH 7.4, 150 mM NaCl, 5 mM MgCl_2_, with 1 mM DTT and 100 μg/ml cycloheximide, 1% (vol/vol) Triton X-100, 25 U/ml Turbo DNase (AM2238, Invitrogen) was dripped onto the plates. Cells were scraped, the lysate was collected to fresh 1.5 ml tube, passed ten times through a 26-gauge needle, cleared by centrifugation at 20,000 x g for 10 min, flash-frozen in liquid nitrogen and stored at -80 °C until further use. For isolation of ribosome-protected RNA fragments, 240 μl of the lysate was digested with 6 μl of RNase I (AM2294, 100 U/μl, Invitrogen) at room temperature with rotation. After 45 min 8 μl of SUPERase-In (20 U/μl, AM2694, Invitrogen) was added to reaction and passed through MicroSpin S-400 HR columns (27-5140-01, GE Healthcare) equilibrated with mammalian polysome buffer. RNA was extracted from the flow-through using Trizol LS (10296-010, Invitrogen) followed by depletion of ribosomal RNA fragments with the RiboZero Kit (MRZH11124, Illumina). Ribosome-protected RNA fragments were loaded onto denaturing 17% urea-PAGE gel (EC-829, National Diagnostics) and gel area ranging from 27 nt to 30 nt, defined by corresponding RNA markers, was cut out. Purified RNA fragments were subjected to library generation using 3′ adapter 4N-RA3, 5′ adapter OR5-4N, RT primer RTP and PCR primers RP1 (forward primer) and RPI1-15 (reverse primers, containing barcodes). Libraries were sequenced on a HiSeq 4000 device (Illumina).

### Polysome Fractionation

Polysome fractionation was performed as previously described (Floor and Doudna, 2016), with minor modifications. Media was removed from 100 mm dish with ∼10^7^ cells and washed with ice-cold ddH_2_O containing 100 μM CHX. All subsequent steps were performed chilled at 4°C or on ice. After ddH_2_O aspiration, cells were incubated for 30 min in 450 μL of hypotonic lysis buffer [0.1x polysome base buffer (PBB), 150 mM KCl, 20 mM Tris-HCl pH 7.4, 15 mM MgCl_2_ in ddH_2_O; with 1% Triton-X 100 and 1x protease inhibitor (4693132001, MilliporeSigma). After confirming >95% free nuclei with a hemocytometer, nuclei were pelleted by centrifugation at 1,800 x g for 5 minutes. Cytoplasmic fraction was separated from mitochondria by centrifugation at 10,000 x g for 5 minutes. 300 μL cytoplasmic lysate was layered atop at 7-45% sucrose gradient (Gradient Master, BioComp, Fredericton, Canada) in 1x PBB. Gradients were ultra-centrifuged at 221,600 x g for 2 hours at 4°C (SW-41Ti rotor, 331362, Beckman Coulter, Brea, CA). Gradients were fractionated (Piston Fractionator, BioComp) into 20 x 300 μL fractions with in-line UV-scanning at 254 nm. Fractions were immediately frozen at -20°C for subsequent RNA extraction.

For comparative polysome profiles, the total input per gradient was normalized by taking the area under the curve (AUC) from the beginning of the SSU peak to the end of the last resolved polysome peak and normalizing the are to 100 artificial units. To normalize for sedimentary variation between gradients, the distance of the SSU, LSU, monosome, 2-, 3-, 4-some peaks across all gradients were averaged, and a simple linear regression was applied with the ‘*ln*’ function in *R* (v 3.5.1) to regress peaks to a mean distance. Raw and processed data and analysis scripts are available in the online electronic notebook.

### Ribosomal molecular dynamics simulations

All molecular dynamics (MD) simulations were performed as previously described (Girodat et al., 2019). In brief, 80S ribosome models were derived from available human Cryo-EM structures with a resolved 18S.1248.m^1^acp^3^Ψ (PDB: 6EKO for E-site tRNA and 6OLE for A/P and P/E tRNA) (Li et al., 2019; Natchiar et al., 2017).

For simulations lacking complete m^1^acp^3^Ψ modifications, the base was converted to each respective precursor. Each system was protonated with the *psfgen* package in VMD 1.9.3, and only e-nitrogen for histidine were protonated (Humphrey et al., 1996). Each system was solvated with a 10Å TIP3P water box with a concentration of 7mM MgCl2 and 100mM KCl using the *solvate* and *autoionize* packages, respectively (Humphrey et al., 1996). All minimizations and MD simulations were performed with *NAMD* 2.1.2 using *CHARMM 36* standard parameters (Best et al., 2012; Denning et al., 2011; Phillips et al., 2005) and modified nucleic acid parameters from (Xu et al., 2016).

Each system underwent a steepest descent minimization of water for 10,000 steps then water and ions for 100,000 steps twice followed by minimization of nucleic acid and protein for 50,000 steps and finally the whole system for 100,000 steps. After minimization all systems were equilibrated to 300 and 350 K for 150 ps. Coordinates of the 350 K equilibration in conjunction with velocities from the 300 K equilibration were used as initial parameters for the MD simulation. Each system was simulated for ∼20ns. RMSD, RMSF, dihedral angel, and distance measurements were performed with VMD 1.9.3 (Humphrey et al., 1996).

### Tandem Mass Tag labeling and liquid-chromatography and tandem mass spectrometry (TMT LC-MS/MS)

HCT116 [WT] and TSR3[KO/Het] cells were grown to 80% confluence in a 100mm dish, washed in PBS, detached, and washed twice in PBS. Cell suspensions were aliquot into technical triplicates and pelleted before snap freezing. One of each HCT116 [WT] and duplicates of HCT116 *TSR3*[KO/Het] clones were processed for LC-MS/MS. Pellets were lysed in 200 μL lysis buffer [4M Guanidine HCl (G4505-500G, Sigma), 40 mM 2-chloroacetamide (22790, Sigma), 10mM tris(2-carboxyethyl) phosphine (C4706-2G, Sigma), 5 mM Ethylenediaminetetraacetic acid (AM9260G, Invitrogen), 1x phosSTOP inhibitor (4906845001, Sigma), 1x cOmplete protease inhibitor cocktail, EDTA free (4693132001, Sigma), 50 mM HEPES (H3784-100G, Sigma)].

The cells were lysed by bead beating using Lysing D matrix tubes (116913100, Cedarlane Laboratories) in a FastPrep 24 instrument (116005500, MP Biomedical) at 8 M/s for 30 seconds, resting 90 seconds, and a second round of bead beating at 8 M/s for 30 seconds. Supernatant was transferred into 1.5 mL lo-bind tubes (022363204, Eppendorf) Protein were solubilized and denatured by heating at 95 °C for 15 minutes with shaking at 1000 rpm in a thermomixer (05-400-205, Fisher Scientific).

All experiments used a 1:1 combination of two different types of carboxylate-functionalized beads, both with a hydrophilic surface (45152105050350 and 65152105050350, Sera-Mag Speed beads, GE Life Sciences).The magnetic particles are an average diameter of 1μm. The beads were stored at 4 °C at a concentration of 10 μg/μL. Magnetic racks used in all experiments were designed and manufactured in-house as previously described (Hughes et al., 2014).

The Sera-Mag SP3 1:1 bead mix was rinsed once with 200 uL HPLC water (270733, Sigma,) and diluted to a final working concentration of 20 μg/μL. To each sample, 200μg of bead mixture was added and mixed by pipetting to generate a homogeneous solution. To induce protein binding to the beads, ethanol (34852, Sigma) was added to achieve a final concentration of 50% (v/v). Bead-protein solutions were mixed to ensure a homogeneous distribution of the beads and incubated for 5 minutes at 24 °C with 1000 rpm shaking in a thermomixer. After incubation, tubes were placed on a magnetic rack for 2 minutes then the supernatant was removed and discarded. The beads were rinsed three times with 200 μL of freshly prepared 80% ethanol in water, and the supernatant was discarded each time. Rinsed beads were reconstituted in aqueous buffer (100μL, 50 mM HEPES pH 8.0) containing a 1:50 (μg:μg) enzyme to protein amount of trypsin/LysC mix (V5071, Promega). Mixtures were incubated for 16 hours at 37°C with 1000 rpm shaking in a thermomixer. The supernatants were recovered using a magnetic rack and transferred to fresh 1.5mL lo-bind Eppendorf tubes.

Tandem Mass Tag (TMT) labels (5 mg per vial, <u>90406</u>, Pierce) were reconstituted in 500 μLof HPLC acetonitrile (34851-4L, Sigma) and frozen at -80 °C. Prior to labeling, TMT labels were removed from the freezer and allowed to equilibrate at room temperature. A standard 13 cell line “supermix” digest mix was prepared as described above and included as an internal standard. Labeling was performed by addition of TMT label in two volumetrically equal steps of to achieve a 2:1 (μg:μg) TMT label to peptide final concentration with 30 minutes incubation at room temperature for each labeling step. Reactions were quenched by addition of an equal volume of 1 M glycine (G8898, Sigma) in water. Labeled peptides were concentrated in a SpeedVac centrifuge (781001010234 + RVT400-115, Labconco) to remove acetonitrile, combined, acidified to a concentration of 1% trifluoroacetic acid (T6508-100mL, Sigma,) and peptides were purified using a C18 SepPak (WAT054960, Waters). The SepPak was conditioned with 2 mL 0.1% trifluoroacetic acid in HPLC grade acetonitrile and equilibrated with 2 mL of 0.1% trifluoroacetic acid in HPLC water. The sample was loaded on the SepPak and rinsed with 3 mL of 0.1% formic acid (, 56302-50ML-F, Sigma) in HPLC water. Peptides were eluted using two 600 uL aliquots of 80% acetonitrile in HPLC water with 0.1% formic acid.

High-pH reversed phase analysis was performed on an Agilent 1100 HPLC system (G1364C, G1315A, G1313A, G1311A). Fractionation was performed on a Kinetix XB C18 column (2.1 x 150 mm, 1.7 μm core shell, 100 Å, Phenomenex, 00F-4498-AN). Elution was performed with a gradient of mobile phase A (water and 0.1% formic acid) to 7% B (acetonitrile and 0.1% formic acid) over 2 minutes, to 25% B over 94 minutes, to 40% over 17 minutes, with final elution (80% B) and equilibration (5% B) using a further 7 minutes all at a flow rate of 450 nL/min. Fractions were collected every minute across the elution window for a total of 48 fractions, which were concatenated to a final set of 12 (e.g. 1 + 13 + 25 + 37 = fraction 1). Fractions were dried in a SpeedVac centrifuge and reconstituted in 0.1% formic prior to MS analysis.

Analysis of TMT labeled peptide fractions was carried out on an Orbitrap Fusion Tribrid MS (IQLAAEGAAPFADBMBCX, Thermo Scientific). Samples were introduced using an Easy-nLC 1000 system (LC120, Thermo Scientific). Trapping columns were packed in 100 μm internal diameter, 360 um outer diameter fused silica capillary (1068150023, Molex) to a length of 25 mm with C18 beads (Reprosil-Pur, Dr. Maisch GmbH, 3 μm particle size, r13.b9.0001). The analytical column was packed with C18 (Reprosil-Pur, Dr. Maisch, 3 μm particle size) to a length of 15 cm in a 100 μm 100 μm internal diameter, 360 um outer diameter fused silica capillary with a laser-pulled electrospray tip. Trapping was carried out for a total volume of 10 μL at a pressure of 400 bar. After trapping, gradient elution of peptides was performed on and heated to 50 **°**C using AgileSLEEVE column ovens (AS1032 + AS1532H, Analytical Sales & Service). Elution was performed with a gradient of mobile phase A (water and 0.1% formic acid) to 8% B (acetonitrile and 0.1% formic acid) over 5 minutes, to 25% B over 88 minutes, to 40% over 20 minutes, with final elution (80% B) and equilibration (5% B) using a further 7 minutes at a flow rate of 375 nL/min.

Data acquisition on the Orbitrap Fusion (control software version 3.1.2412.17) was carried out using a data-dependent method in positive ion mode. Survey scans covering the mass range of 400 – 1200 were acquired in profile mode at a resolution of 120,000 (at *m/z* 200), with quadrupole isolation enabled, an S-Lens RF Level of 60%, a maximum fill time of 120 ms, an automatic gain control (AGC) target value of 4e5, and one microscan. For MS2 scan triggering, monoisotopic precursor selection was enabled for peptides, charge state filtering was limited to 2 – 4, undetermined charge states were included, and dynamic exclusion of masses selected one time was enabled for 15 seconds with a tolerance of 20 ppm. HCD fragmentation with a maximum fill time of 20 milliseconds, quadrupole isolation, an isolation window of 1.4 *m/z*, isolation offset off, a fixed collision energy of 40%, injection for all available parallelizable time turned OFF, and an AGC target value of 1.2e5 was performed. The Orbitrap was used as the detector in normal scan range mode with a resolution of 50,000 at 200 *m/z* and the first mass was 120 m/z. Centroided data was collected with one microscan. The total allowable cycle time was set to 3 seconds.

### LC-MS/MS data analysis

Peptide detection and protein inference was performed using *Proteome Discoverer* (v 2.4). Searches were performed using *Sequest HT* and the human proteome from Uniprot plus the CRAP-ome. Tryptic cleavages with 2-6 missed cleavages and peptide length of 6-144 was used. The precursor mass tolerance was 10 ppm, fragment mass tolerance was 0.05 Da and monoisotopic masses were used. Only b- and y-type ions were included in the search. Oxidation was allowed as a dynamic modification and the protein terminus was allowed to have acetylation, Met-loss, or Met-loss+acetylation as dynamic modifications. TMT reagents (+229.163 Da) were set as static modifications on the peptide terminus and lysine and carbamidomethyl was assigned as a static modification of cystine. Percolator was used for scoring with an FDR of 0.01, concatenated Target/Decoy Selection, and validation based on the q-Value. The maximum Delta Cn was 0.05 and the maximum rank was 0. Unique + Razor peptides were used for quantification. *R* (v 3.5.1) was used for data analysis. Peptides were filtered for uniqueness and values were median normalized, and proteins containing at least two unique peptides were included in downstream differential protein expression analysis. The technically replicated samples were merged (mean) and statistical protein expression analysis was performed in *R.*

### Statistics

Statistical analysis was performed in *R* (v 3.5.1). Differences in variant allele frequency (VAF) between tumor and normal patient samples was two-tailed, paired Student’s T-test with degrees of freedom one less than reported *n.* Bonferonni multiple-testing correction was applied when screening for changes across 18S and 28S nucleotides. Error bars on boxplots are quantiles. Differential gene expression and translation was tested with *DEseq2* (Love et al., 2014) with Benjamini-Hochberg multiple testing correction at an alpha of 0.05. Multi-group comparisons between HCT116 WT[1],[2],[3] and *TSR3*[KO 1],[KO 3],[Het2] ribo-seq were performed with one-way ANNOVA, followed by Tukey’s Honestly Significant Difference (HSD) test if indicated. Genotypic differences between biological triplicates of WT and *TSR3*[KO] cells were tested with Student’s T-test with 2 degrees of freedom and Benjamini-Hochberg multiple testing correction at an alpha of 0.05 where indicated.

