## Supplementary figures and images for "Loss of m^1^acp^3^Ψ ribosomal RNA modification is a major feature of cancer"

### Supplementary Figures

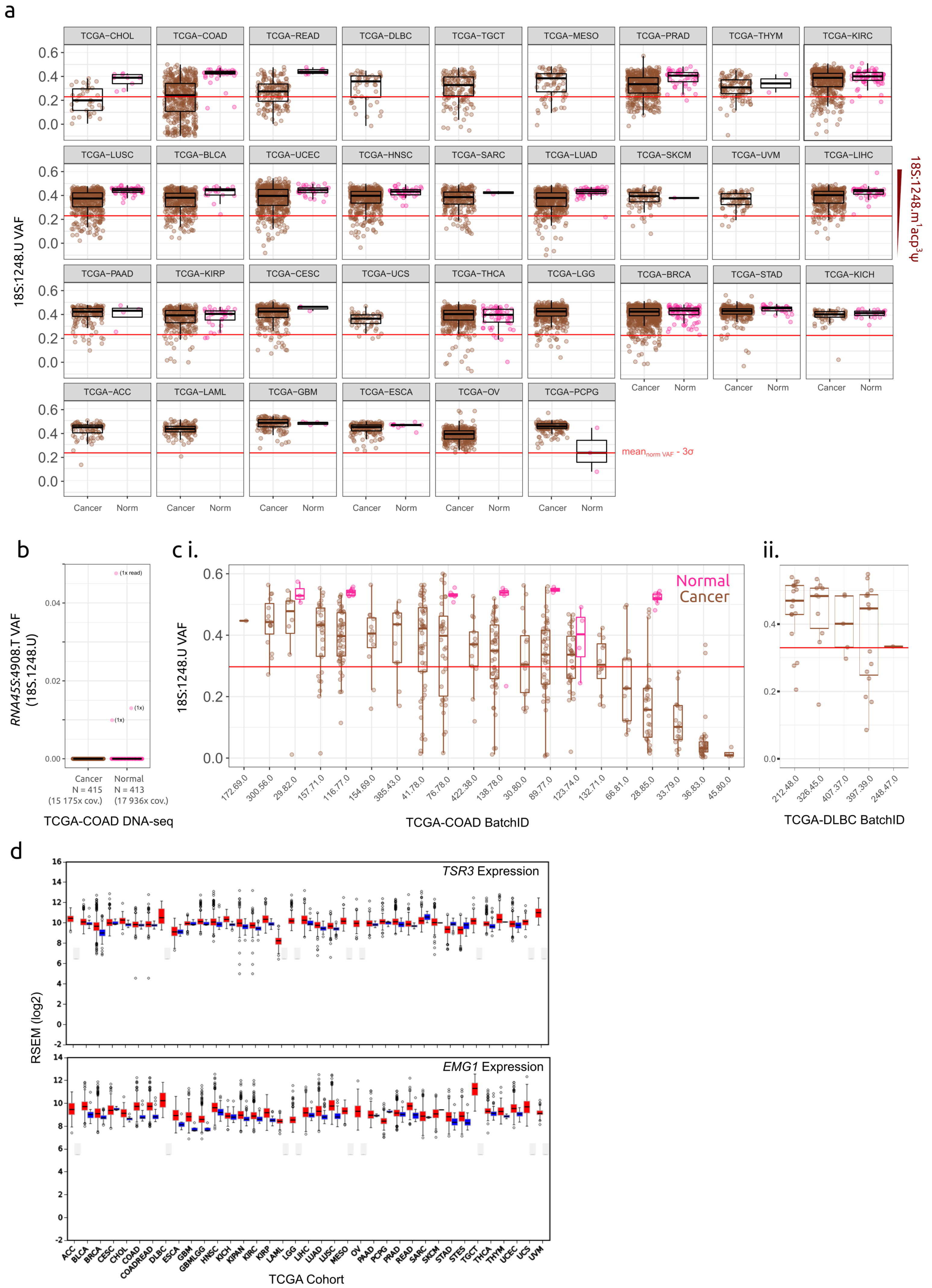

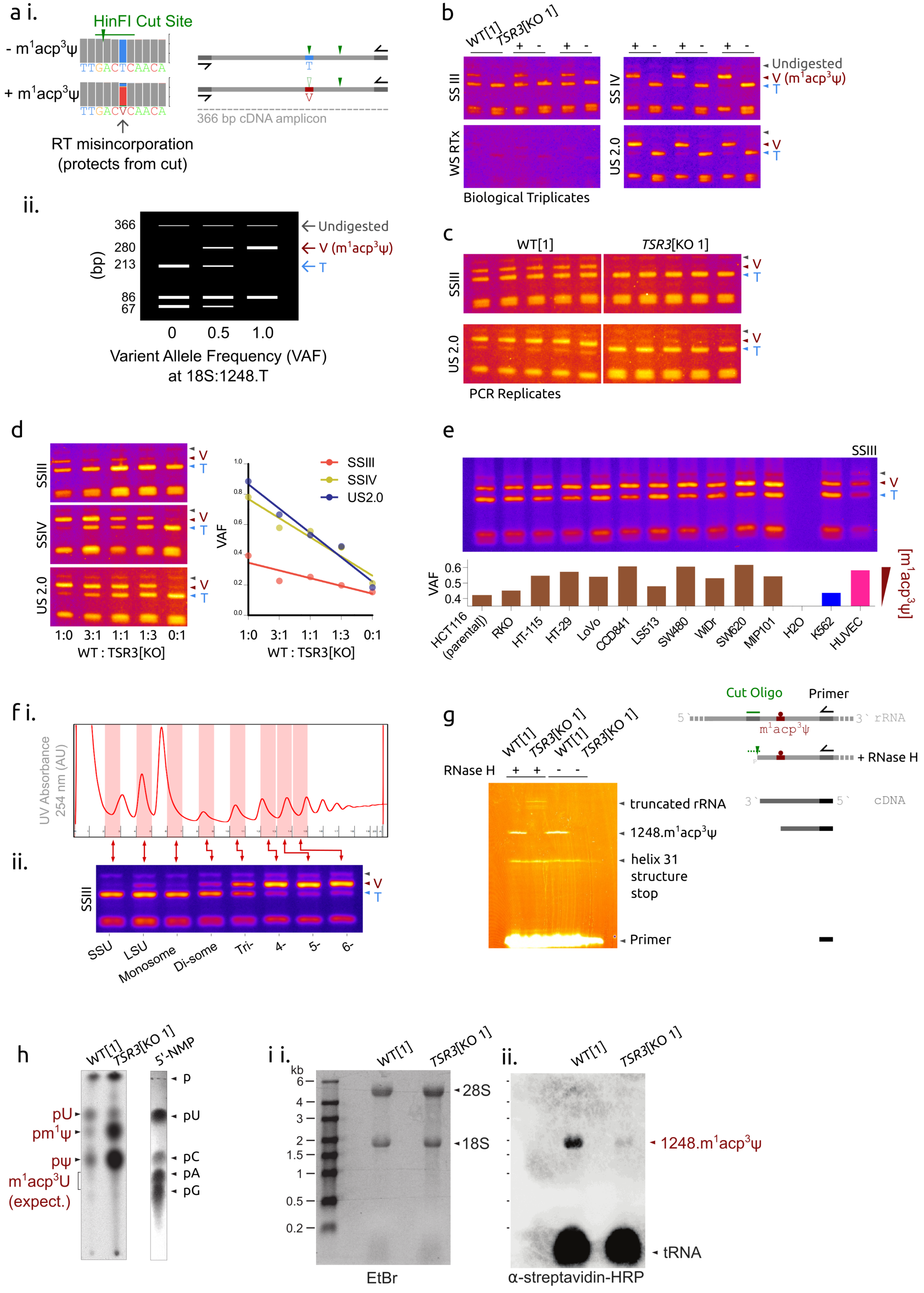

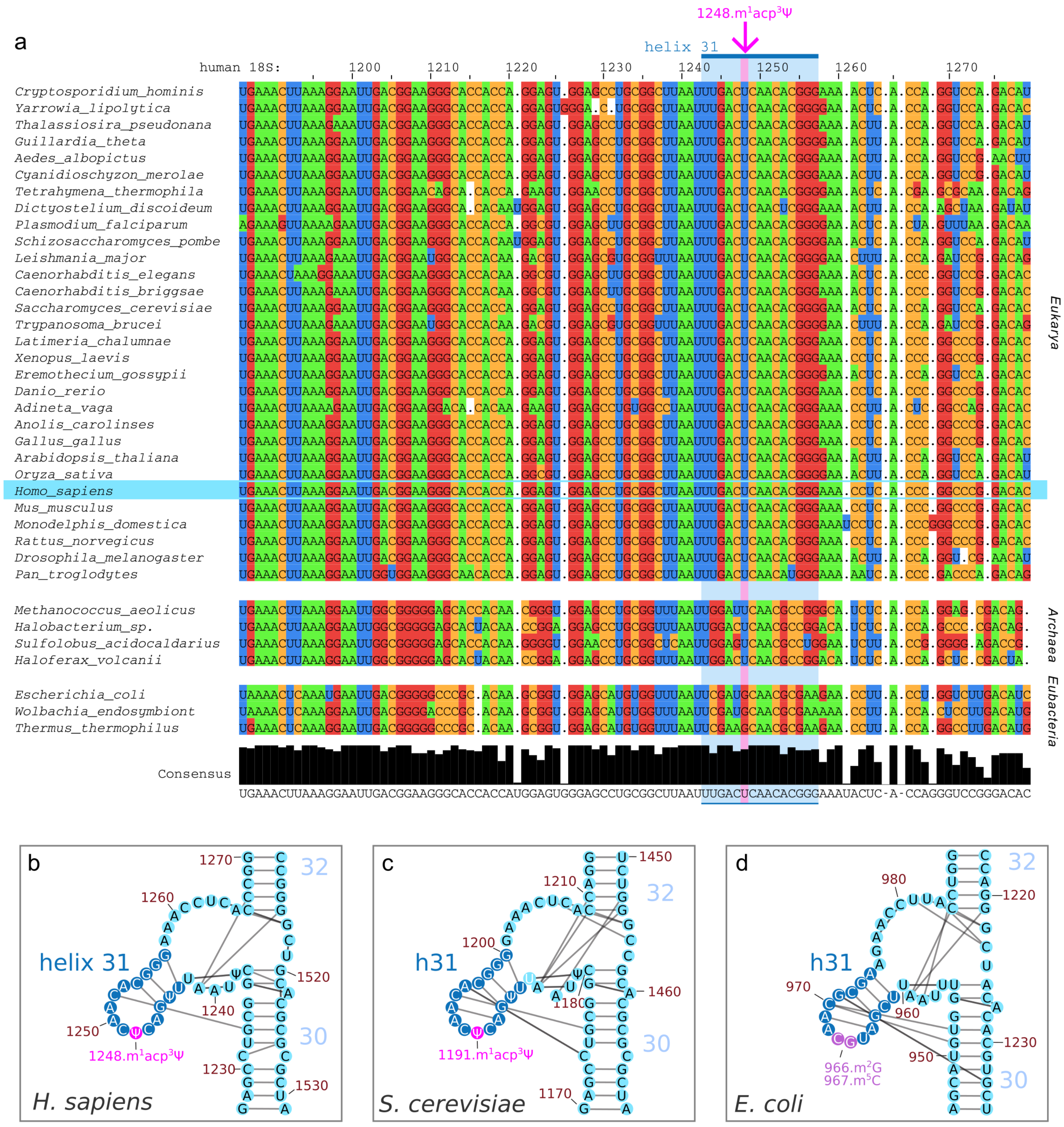

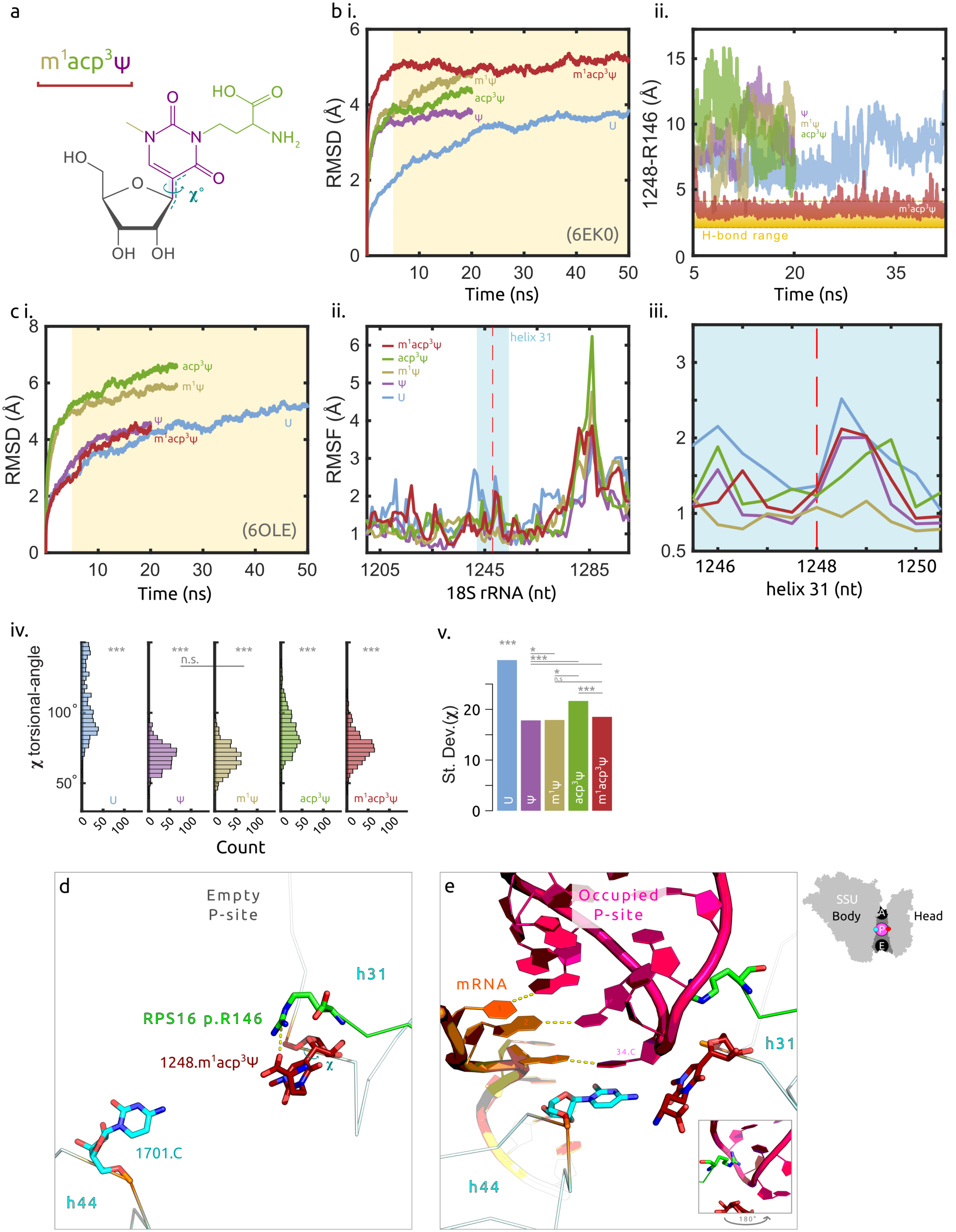

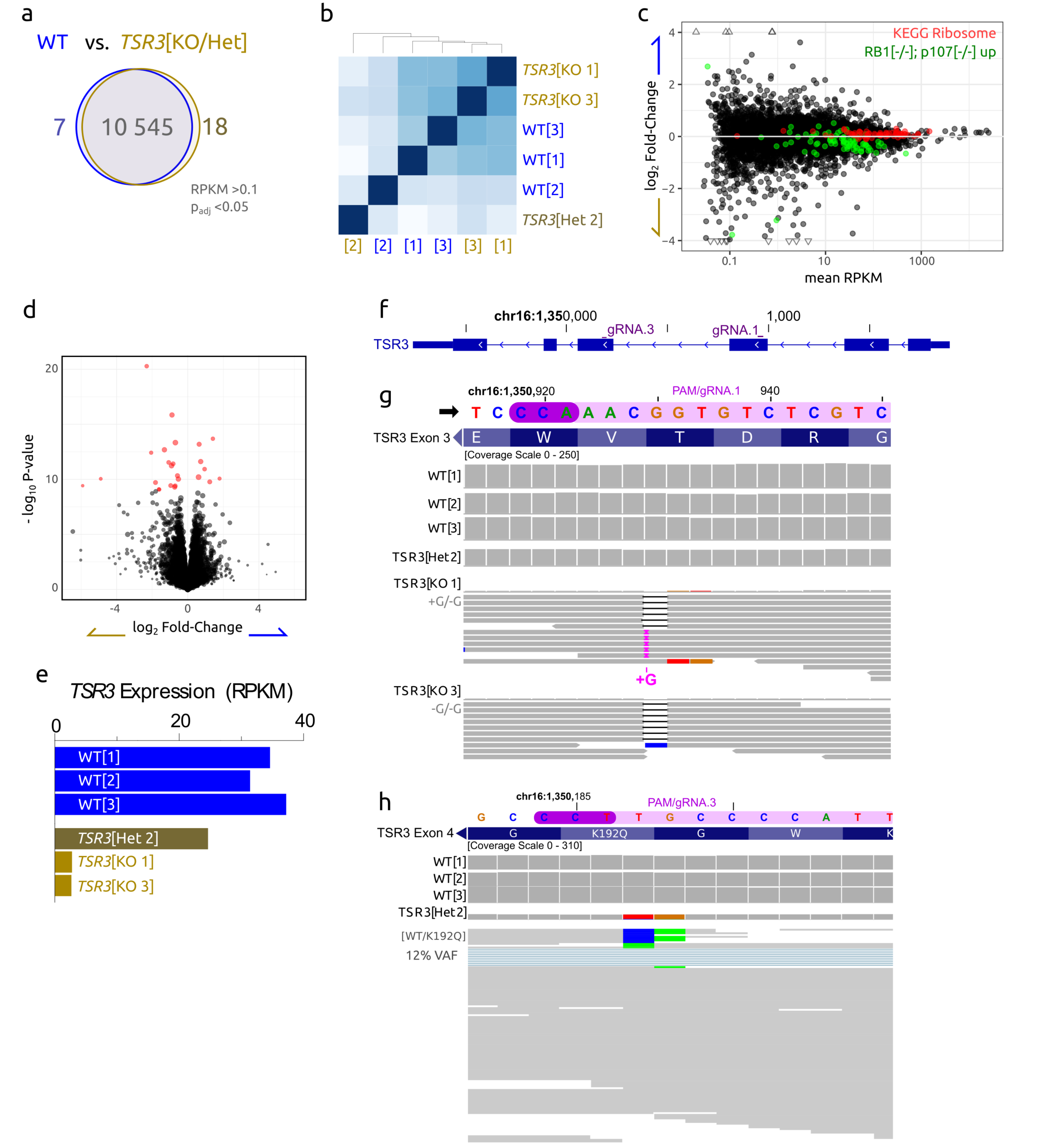

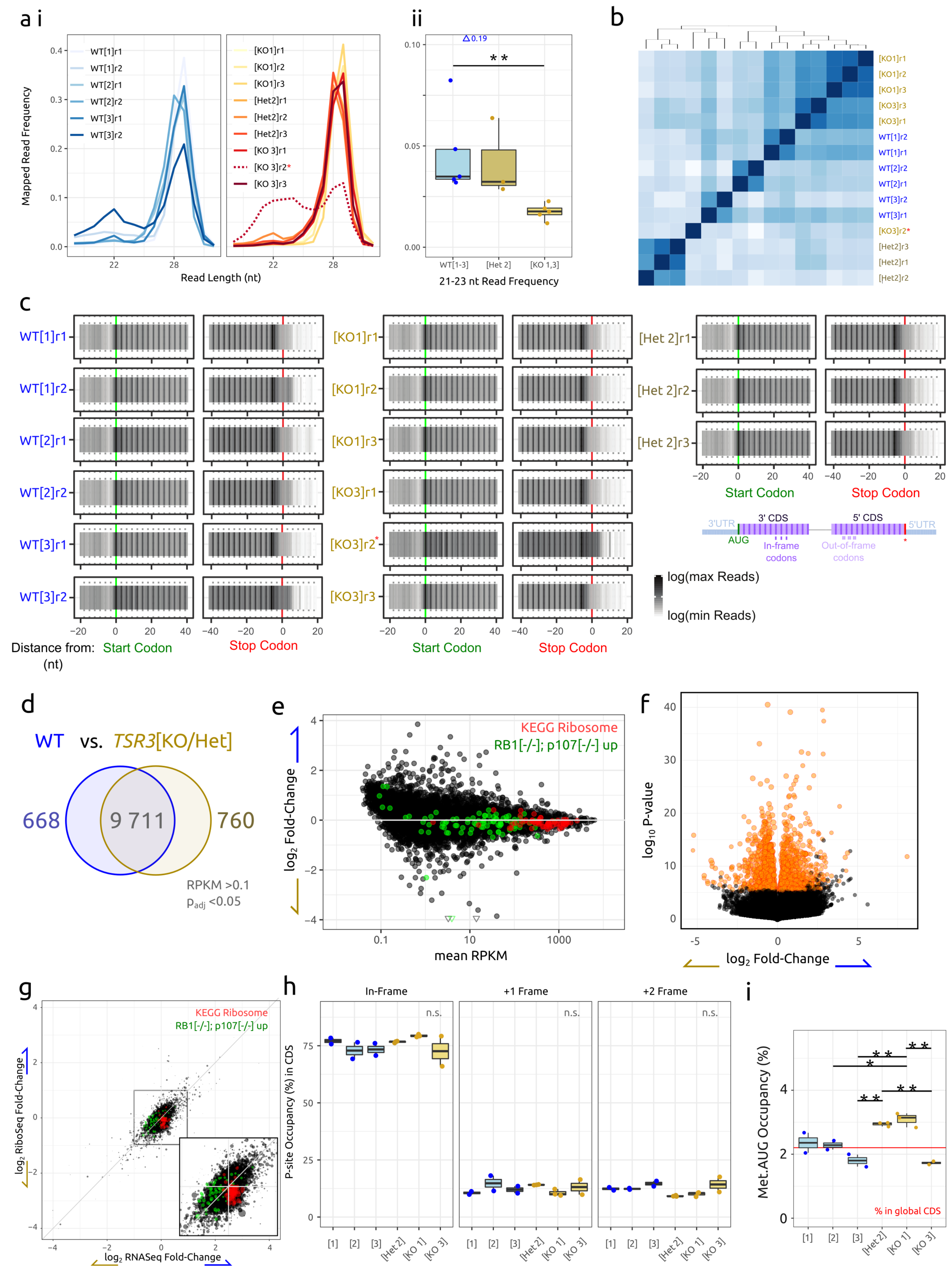

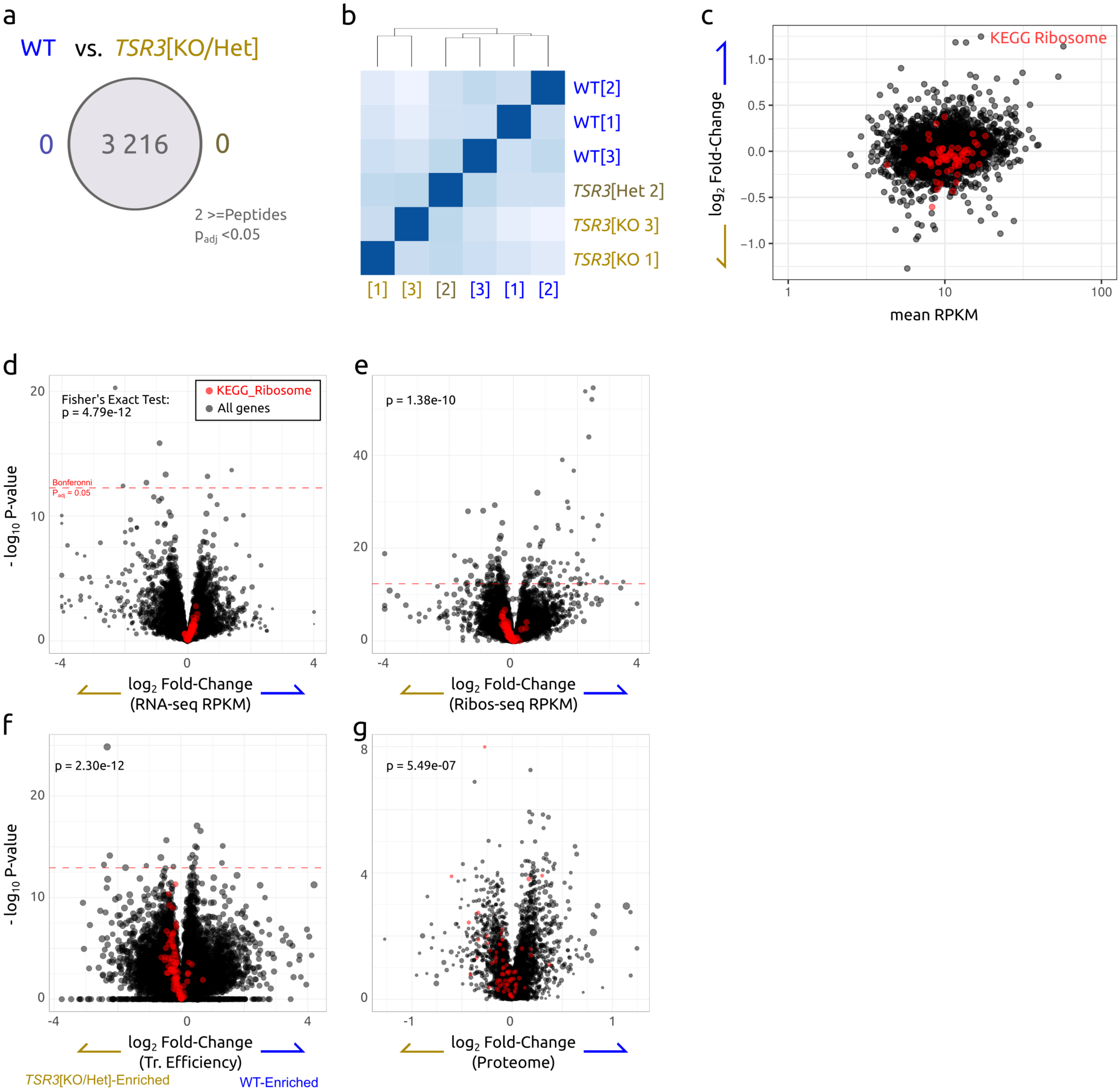

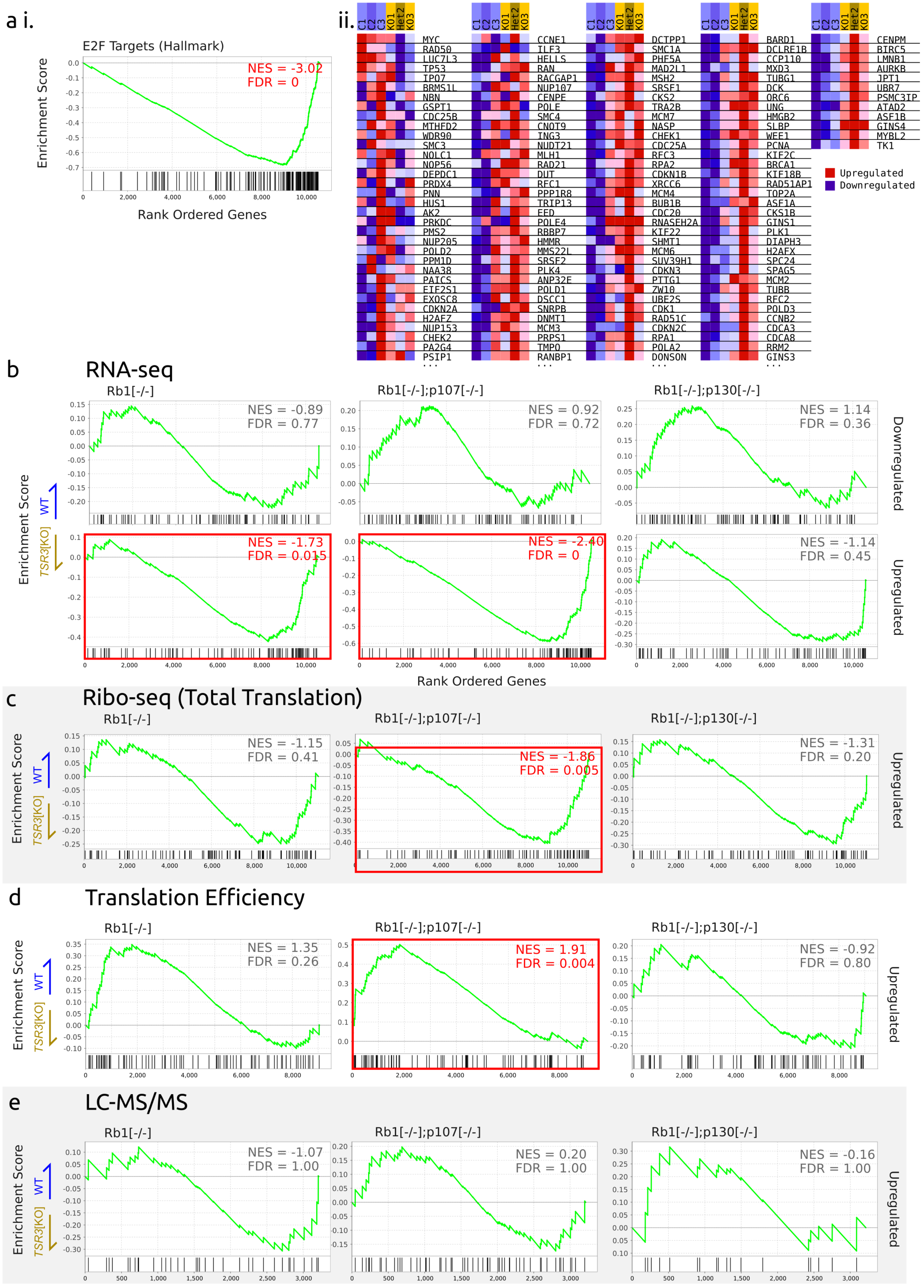

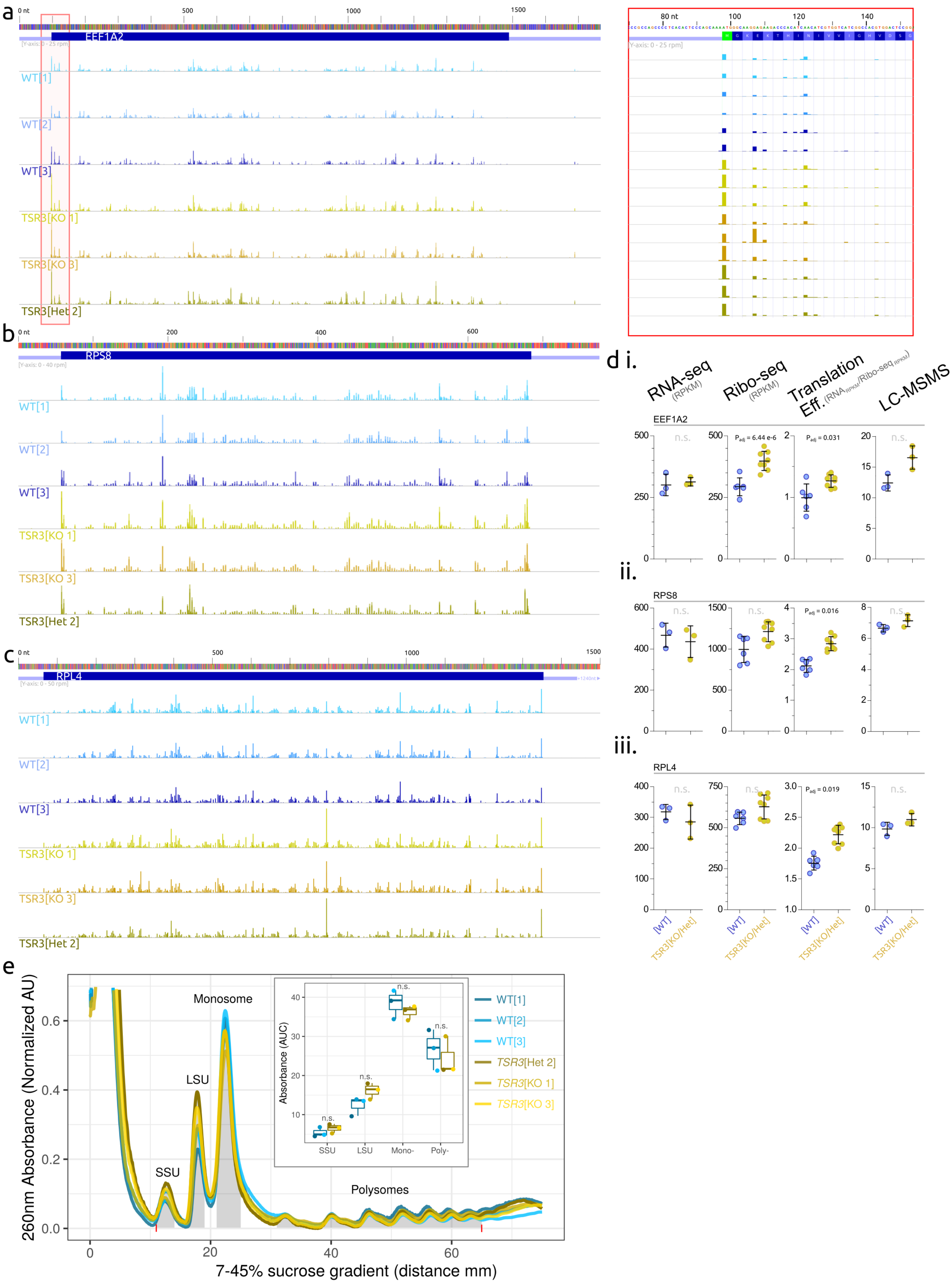
